# MaxEnt-DTD: Maximum-Entropy Estimation of Diffusion Tensor Distribution for Fiber Orientation and Microstructure Characterization

**DOI:** 10.64898/2026.06.19.733471

**Authors:** Yiang Pan, Yuanjing Feng, Jianzhong He, William Consagra, Carl-Fredrik Westin, Yogesh Rathi, Lipeng Ning

## Abstract

Diffusion MRI (dMRI) enables noninvasive characterization of white-matter fiber orientations and tissue microstructure, but widely used approaches, such as constrained spherical deconvolution (CSD) and parametric multicompartment models, typically address these features separately. The diffusion tensor distribution (DTD) framework jointly represents fiber orientation and microstructure, but estimating DTD from finite, noisy measurements is severely ill-posed. Existing inversion methods either rely on nonnegativity constrained basis representations, which are challenging to sale to high-dimensional and high-resolution distributions, or use sampling-based approaches with limited reliability. We propose MaxEnt-DTD, a maximum-entropy algorithm for DTD estimation from finite and noisy dMRI data. By deriving the Lagrange dual formulation, we reformulate a constrained infinite-dimensional optimization problem into a finite-dimensional unconstrained convex optimization problem, substantially reducing the parameter space and enabling tractable whole-brain DTD estimation. We evaluate MaxEnt-DTD using both synthetic and *in vivo* data from the Human Connectome Project protocol and a second dataset using advanced B-tensor diffusion encoding. We compare MaxEnt-DTD-derived fiber orientation distributions with results from CSD and Monte-Carlo inversion methods, and assess fiber-specific microstructure measures and rotation-invariant metrics based on the cumulants of DTD. The results demonstrate that MaxEnt-DTD provides a reliable and efficient framework for joint fiber-orientation and microstructure analysis in dMRI.

## 1. Introduction

Diffusion magnetic resonance imaging (dMRI) is a non-invasive imaging modality for characterizing brain tissue microstructure using pulsed magnetic field gradients (Le Bihan et al., 1986; Basser et al., 1994a; Callaghan, 2011; Novikov et al., 2019). dMRI signals are sensitive to water diffusion at the micrometer scale and provide information about tissue microstructure. Several dMRI analysis techniques have been developed to derive imaging markers for disease diagnosis, neurosurgical planning, and the investigation of neurodegeneration (Essayed et al., 2017; Novikov et al., 2019). These methods have often relied on separate techniques to address two distinct objectives: characterization of white-matter fiber orientations and quantification of tissue microstructure (Özarslan et al., 2013; Ning et al., 2015; Jelescu and Budde, 2017; Novikov et al., 2019).

Gradient direction-dependent dMRI signals are sensitive to fiber orientations that can resolve crossing white-matter bundles to compute tractography and characterize brain connectomics (Tuch et al., 2002; Sporns et al., 2005; Wedeen et al., 2008). Constrained spherical deconvolution (CSD) is one of the most widely used methods for fiber orientation distribution (FOD) estimation (Tournier et al., 2007; Descoteaux et al., 2008), which has been extended several variations to characterize mixtures of different tissue types (Jeurissen et al., 2014, 2019; Raffelt et al., 2017) and using advanced diffusion encoding waveforms (Jeurissen and Szczepankiewicz, 2021; Karan et al., 2022). Despite these advances, the CSD framework does not recover the microstructure across fiber bundles and uses a fixed microstructure kernel as the fiber response function, which ignores heterogeneity between crossing fiber bundles (Christiaens et al., 2020; De Santis et al., 2014).

In addition to fiber orientations, dMRI signals measured with different gradient strengths or b-values are also sensitive microscopic cellular arrangement (Beaulieu, 2002; Basser et al., 1994b). Standard dMRI modeling techniques, such as diffusion tensor imaging (DTI) (Basser and Pierpaoli, 1996) and diffusion kurtosis imaging (DKI) (Jensen et al., 2005), estimate the cumulants of water diffusion displacements to derive quantitative metrics such as mean diffusivity (MD), fractional anisotropy (FA) (Basser and Pierpaoli, 1996) and mean kurtosis (MK) to characterize tissue microstructure (Novikov et al., 2019; Alexander et al., 2019). More advanced methods have been developed to extend the finite moments to the probability distribution of water displacements, i.e., the ensemble average propagator, to simultaneously resolve crossing fibers and provide microstructure measures (Wedeen et al., 2008; Ning et al., 2015; Özarslan et al., 2013). But these methods do not provide information on the microstructure of specific fiber bundles and have limited biophysical interpretations.

Parametric dMRI models have been developed to quantify microstructural tissue properties. Most methods decompose dMRI signals into a linear mixtures of signals from multiple tissue compartments which have different analytical expressions derived from biophysical models. The examples include the widely used NODDI (Zhang et al., 2012), SANDI (Palombo et al., 2020) and the Standard Model framework (Novikov et al., 2018). While these parametric models provide specific tissue microstructure measures, they often rely on strict assumptions that may bias estimation when the underlying microstructure deviates from the assumed form(Lampinen et al., 2017; Jelescu et al., 2016). Moreover, the parametric models for white matter are usually developed for a single dominant fiber direction. Modeling of dMRI signals for multiple crossing fiber bundles requires a higher-level mixture of these models for different directions, which further increases the complexity of solving the inversion problem. Existing methods, such as (Consagra et al., 2025), typically assume of a fixed parametric microstructure model and pre-specified fiber orientations.

The diffusion tensor distribution (DTD) is a general framework for characterizing tissue heterogeneity without the need for a pre-specified number of compartments or a single shared response kernel (Jian et al., 2007; Westin et al., 2016; Topgaard, 2019; Lasič et al., 2014a; Lampinen et al., 2017). In this framework, tissue microstructure is characterized by a continuous distribution of microscopic diffusion tensors where each tensor characterizes a local tissue enviorment within a voxel. These microscopic tensors may have distinct orientation and diffusivity, representing hetergenouse tissue microstructure. Several theories and techniques have been developed to estimate the cumulants of DTD (Westin et al., 2016; Ning et al., 2021a). Tensor-valued diffusion encoding waveforms are needed to estimate higher-order cumulants that can provide microstructure measures, such as microscopic anisotropy, that cannot be provided by standard linear encoding methods. But the cumulant-based methods do not directly provide information to resolve crossing fiber bundles.

Estimating the DTD beyond the cumulants using finite and noisy dMRI data is an extremely ill-posed problem. Basis-representation approaches characterize the signal using a large dictionary matrix with each column representing signals from a microscopic tensor in the sample space, resulting in a large matrix needed for high-dimensional and high-resolution DTD estimation. Standard inversion methods, such as non-negative least squares (NNLS), suffer from strong coupling between basis signals and limited spectral resolution that degrades with increasing dimensionality(de Almeida Martins and Topgaard, 2016). Monte Carlo inversion(Topgaard, 2017; de Almeida Martins and Topgaard, 2018; Topgaard, 2019; Magdoom et al., 2021) avoids fixed grids but introduces stochastic variability that may have limited reproducibility and increased sensitivity to noise (Magdoom et al., 2021; Reymbaut et al., 2021b).

In this paper, we introduce a new algorithm to estimate the DTD motivated by the maximum-entropy (MaxEnt) principle for the moment problem (Jaynes, 1957). We extend the MaxEnt framework introduced in our previous work (Ning, 2023a) for a different inverse problem in a lower-dimensional space to high-dimensional DTD estimation. The present formulation introduces a scalable approach to estimate DTD, which can simultaneously resolve crossing fiber bundles and provide fiber-bundle-specific microstructure measures. The main contributions of this paper are:

1. A maximum-entropy formulation for DTD estimation that solves a finite-dimensional unconstrained convex optimization problem with a unique global optimum. We introduce a discrete optimization algorithm with adaptive sampling in the fiber-orientation and microstructure spaces to improve computational efficiency and estimation performance.
2. A unified framework that jointly estimates fiber orientation distributions, fiber-specific microstructure distribution, and orientation-invariant microstructure measures.
3. A systematic evaluation of the proposed MaxEnt-DTD method for angular resolution and microstructure estimation, with comparisons to state-of-the-art approaches, including CSD for FOD estimation, Monte Carlo (MC) inversion for DTD estimation, and cumulant-based DTD analysis methods.
4. A comprehensive evaluation using synthetic and *in vivo* dMRI data acquired with conventional multi-shell linear encoding and tensor-valued encoding, providing quantitative benchmarks and practical guidance for applying the proposed method.

## 2. Methodology

### 2.1. Signal model and parameterization

In the DTD framework, each voxel is represented as a heterogeneous collection of microscopic environments, where water diffusion within each environment is characterized by a positive-definite diffusion tensor **D** (Westin et al., 2016). The size, shape and orientation of the micrometer-scale microenvioments are characterized by the eigenvalues and eigenvectors of the tensor **D**. Each millimeter-scale voxel contains contributions from a multitude of such microenvironments, and these contributions are modeled by a continuous distribution of diffusion tensors, referred to as the diffusion tensor distribution (DTD), *P* (**D**) (Jian et al., 2007; Westin et al., 2016; Topgaard, 2017, 2019). Accordingly, the dMRI measurement is represented as

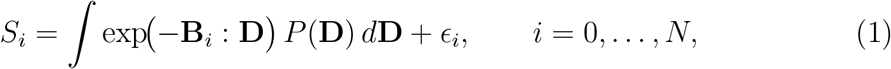

where *S*_*i*_ represents the dMRI measurement related to signal, **B**_*i*_ is the *i*-th diffusion encoding tensor, **B**_*i*_ : **D** denotes the Frobenius inner product ∑_*m,n*_[**B**_*i*_]_*mn*_[**D**]_*mn*_ (Karan et al., 2022), and *N* is the total number of measurements and *ϵ*_*i*_ represents the Gaussian measurement noise which is valid with relatively high SNR (Sijbers et al., 1998; Gudbjartsson and Patz, 1995). Different from the formulation in (Reymbaut, 2020), the *S*_*i*_ is not the normalized dMRI signal and the density *P* (**D**) is not restricted to a probability density function whose total mass, i.e., ∫ *P* (**D**)*d***D**, represents the noise-free baseline signal. This representation does not reduce any information for microstructure estimation but simplifies the algorithm to solve the inverse problem, which will be explained in the following sections.

#### 2.1.1. Axisymmetric parameterization

To reduce the dimensionality of the inverse problem while still providing relevant metrics, we assume the microscopic diffusion tensor **D** has axisymmetric structure as in (Eriksson et al., 2015; Lasič et al., 2014b). Specifically the tensor **D** is parameterized by four parameters

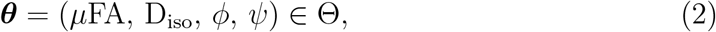

where *µ*FA ∈ [0, 1) quantifies the microscopic anisotropy, i.e., the “shape”, D_iso_ characterizes the microscopic mean diffusivity, i.e., the “size”, and (*ϕ, ψ*) ∈ [0, *π*] × [0, 2*π*) parameterize orientation of the dominant eigenvector. Depending on the context, the parameter may also be represented as (*µ*FA, D_iso_, **u**) with **u** = **u**(*ϕ, ψ*) ∈ S^2^.

The axisymmetric diffusion tensor is constructed as

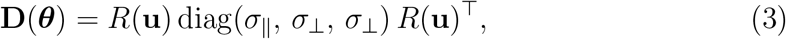

where *R*(**u**) is the rotation matrix aligning the principal axis with **u** with mapping from (*µ*FA, D_iso_) to eigenvalues is given by

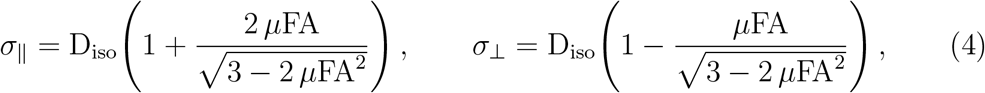

which guarantees *σ*_⊥_ ≥ 0, *σ*_∥_ ≥ *σ*_⊥_, and *σ*_∥_ + 2*σ*_⊥_ = 3 D_iso_ for all admissible (*µ*FA, D_iso_).

Under this parameterization, the signal model in Eq. (1) can be written as

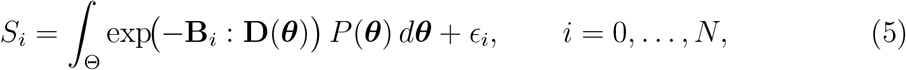

where *P* (***θ***) denotes a distribution in the four-dimensional space Θ. The estimation algorithm is introduced in the next section.

### 2.2. Maximum-entropy estimation of the DTD

#### 2.2.1. Problem formulation

The problem for estimating the continuous distribution *P* (*θ*) using finite measurements is closely related to the moment problem in probability theory where the goal is to determine a probability distribution from its finite sequence of moments. The maximum-entropy (MaxEnt) principle is a classical approach to solve the inverse problem which has led to several well-known theories in the field(Jaynes, 1957; Shannon, 1948; Jaynes, 1982). The principle of maximum entropy has been generalized beyond probability density functions and has been applied in many research fields (Skilling and Bryan, 1984; Phillips et al., 2006; Mohammad-Djafari, 2015). It has been applied in a closely related field to estimate the relaxation-diffusion correlation spectroscopy in (Batou and Soize, 2013; Ning, 2023a).

Below is the proposed more general problem formulation to estimate the DTD:

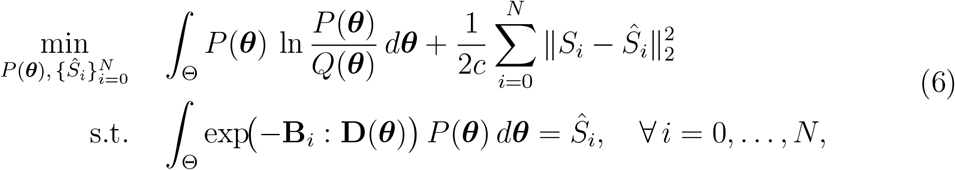

where *Ŝ*_*i*_ represents the estimated noise-free signal, and *c >* 0 is a constant parameter that scales according to the noise level. Without loss of generality, we assume *B*_0_ = 0 and *Ŝ*_0_ is the estimated baseline signal which is the total mass of *P* (***θ***). The first term of the objective function in (6) represents the Kullback-Leibler (KL) divergence between *P* (***θ***) and a known prior density function *Q*(***θ***). When *Q*(***θ***) is a uniform function, then the first term reduces to the negative differential entropy of *P* (***θ***). Then the objective function reduces to the maximum entropy formulation considered in (Batou and Soize, 2013; Ning, 2023a). The more general formulation in (6) makes it possible to integrate prior information to further improve the estimation results. In addition to serving as a prior distribution, a nonuniform *Q*(***θ***) may be necessary when estimating the DTD in different parameter spaces as shown in (Ning, 2023b).

### 2.3. The dual formulation

The problem in (6) is a convex optimization problem with linear constraints. The optimizing variable *P* (*θ*) is a nonnegative function on a compact space Θ. Standard methods, as in (Chouzenoux et al., 2010), use numerical methods to directly solve (6). Similar to standard basis representation techniques, this method requires optimizing a high-dimensional variable, which is computationally inefficient when the problem needs to be solved across all voxels of the whole brain.

Following (Ning, 2023a), we introduce an efficient method to solve (6) by solving the following dual problem

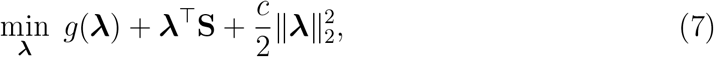

where **S** = [*S*_0_; *S*_1_; … ; *S*_*N*_ ] denote the measured signal vector and

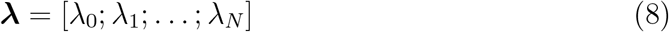

denotes the vector of Lagrange multipliers and

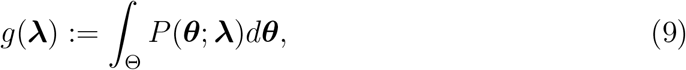

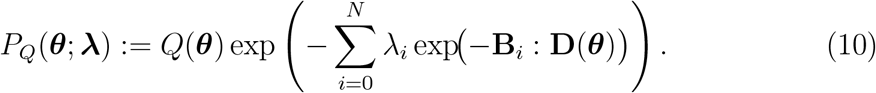

Compared to the primal problem, the dual formulation (7) is a strictly convex optimization problem over a finite dimensional variable ***λ***, ensuring a unique global optimal solution. Let 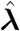 denote optimal solution to (7), then the optimal solution to (6) is given by 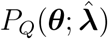. Let *P* (***θ***; ***λ***) denote the special case of *P*_*Q*_(***θ***; ***λ***) when *Q*(***θ***) is the identity function. Then *P* (***θ***; ***λ***) has the same formulation as the standard MaxEnt solution as in (Jaynes, 1957).

We highlight several differences between the proposed dual formulation (7) and related early work. The MaxEnt expression has been used in (Heaton, 2005) for NMR analysis which solves a problem of the following form

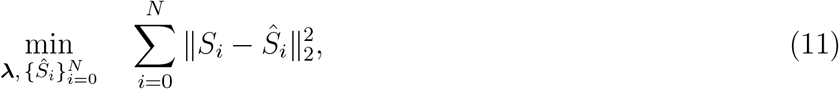

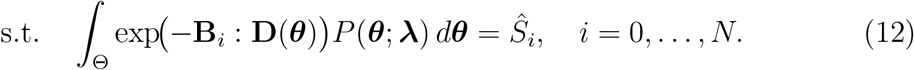

The above optimization problem is not convex and may lead to local optimal solutions. Another closely related method was introduced in (Batou and Soize, 2013) that solves the MaxEnt problem without considering measurement noise and the dual formulation, which may lead to unreliable solutions for noisy dMRI data. Our previous work (Ning, 2023a) has introduced a more general MaxEnt formulation by incorporating measurement noise and the dual formulation. (7) further generalizes the method as in (Ning, 2023a) with an additional prior and also applies the method to estimate DTD for the first time.

### 2.4. Fiber orientation and microstructure estimation

For a given *P* (***θ***), let 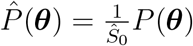 denote the normalized DTD function, with *Ŝ*_0_ denoting the estimated baseline signal which represents the total mass of *P* (***θ***). Several measures can be calculated from the normalized DTD to estimate the fiber orientation and microstructure.

#### Fiber orientation distribution (FOD)

Integrating out the spectral dimensions yields the orientation density on S^2^:

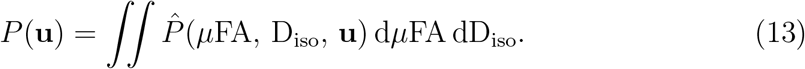

Note that the total mass *P* (**u**) is not normalized by the estimated baseline signal *Ŝ*_0_. *P* (**u**) can be further normalized by 1*/Ŝ*_0_ to adjust inter-voxel differences in signal intensity. Peaks in *P* (**u**) correspond to distinct fiber populations within the voxel and can be compared with fiber orientation distributions obtained from CSD(Tournier et al., 2007; Jeurissen et al., 2014).

However, unlike CSD-derived fiber ODFs, which assume a spatially invariant response function shared across all orientations within a voxel, *P* (**u**) arises as a model-free marginalization of the full DTD and naturally accommodates orientation-dependent microstructural variation. This is a fundamental distinction: while CSD assumes that all fiber populations within a voxel share a common signal response, the MaxEnt-DTD framework permits each orientation to exhibit its own microstructural composition.

#### Orientation-specific microstructure

For a given fiber orientation **u**, the conditional distribution

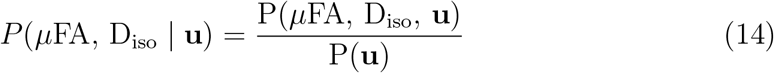

characterizes the size-shape composition along that specific fiber direction. The total number of Peaks in this conditional density funciton indicates the number of microstructural components that may be related to intra-axonal or extra-axonal signal contributions within a single fiber population (Reymbaut et al., 2021a).

#### Rotation-invariant microstructure measures

A family of microstructural measures has been developed based on the cumulant or the moment tensors of the DTD (Westin et al., 2016). These moment tensors can be numerically computed from the estimated DTD functions as

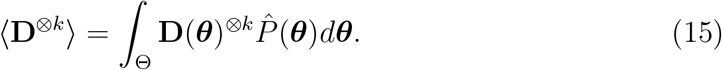

For example, the mean tensor ⟨**D**⟩ can be used to compute the *FA* and mean diffusivity *MD*. The second-order cumulant

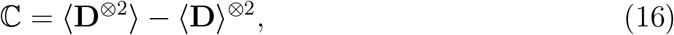

which provides information to compute the mean kurtosis (MK) and more advanced bulk kurtosis (MK_I_) and shear kurtosis (MK_A_).

We note that cumulant expansion-based methods as in (Westin et al., 2016; Ning et al., 2021b; Coelho et al., 2026) usually assume a fixed-order expansion while the MaxEnt approach does not require such constraints. Several theories have been established related to the identifiability of the cumulants. For example the full second-order moment tensor consists of 21 unknown variables that cannot be estimated using linear diffusion encoding methods. However, under the assumption of microscopic axisymmetry, the microstructure diffusion tensor only consists of 4 variables instead of the 6 variables in the full tensor. The corresponding second-order moment for axisymmetric tensors consists of only 10 variables as the results generated by (15) with *k* = 2. On the other hand, standard linear diffusion encoding methods can estimate 15 variables of the second-order diffusion tensor (Westin et al., 2016). Thus, the generative approach in (15) may still provide results to estimate useful information from the second-order moment based on linear diffusion encoding. But the assumption of axisymmetric tensors is a limitation to compute all the rotational invariant metrics based on the full cumulant expansion (Coelho et al., 2026).

### 2.5. Numerical algorithm and adaptive sampling

For numerical implementation, the continuous DTD is discretized on a grid of *M* atoms 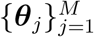. The forward model in Eq. (5) becomes the linear system

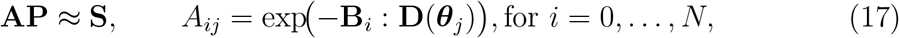

where **S** is the vector of measured signals, **P** ∈ ℝ^*M*^ is the descrete density vector, and **A** is the kernel matrix. Substituting the discrete grid into the dual problem of Eq. (7) with

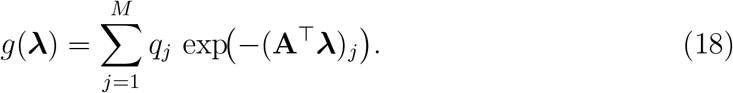

The Hessian of *F* is positive definite (Ning, 2023a), so *F* is strictly convex with a unique global minimizer. Throughout this work, we will consider a uniform prior and impose *q*_*j*_ = 1. The algorithm is generalizable to situations when meaningful priors are available.

A key computational advantage of this formulation is that the optimization is performed in the (*N* + 1)-dimensional dual space rather than the *M* -dimensional primal space, with *N* ≪ *M* in practice. Each evaluation of *F* and its gradient traverses all *M* atoms to compute *g*(***λ***), giving a per-iteration complexity of O(*NM*). We minimize *F* using a truncated Newton method with conjugate-gradient (CG) inner iterations. At each Newton step, the search direction **d** is obtained by approximately solving the Newton system **H d** = −∇*F* via preconditioned CG, where the Hessian-vector products **Hv** are computed without explicitly forming **H**, and a diagonal Jacobi preconditioner **M**^−1^ = diag(**H**)^−1^ accelerates convergence. The CG iterations terminate when the residual falls below a tolerance *ϵ*_CG_ or a maximum iteration count is reached. The Newton step is safeguarded by clamping the largest component of **d** to prevent numerical overflow in the exponential terms. The entire procedure is implemented in PyTorch and executed on GPU, with all voxels processed in parallel batches, enabling whole-brain DTD estimation within a clinically practical timeframe.

The computational results depend on the sampling of the grid points. Although a large number of samples helps improve the resolution of the estimated density, it also increases computational cost. To mitigate this limitation, we introduce an adaptive sampling method based on the assumption that the density functions have sparse support in the angular directions. Specifically, we assume that

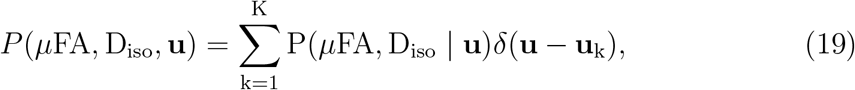

where *δ*(·) represents the Dirac delta function. In the above formulation, the difference between the volume fractions of between fiber bundles is represented by the difference in the conditional density functions.

The first step estimates the fiber orientations by solving (7) using a discrete basis with *M*_1_ = *M*_lr_*M*_**u**_, where *M*_lr_ represents the number of low-resolution samples in the *µ*FA − D_iso_ plane and *M*_**u**_ denotes the number of samples on the unit sphere used to improve angular resolution. In this work, we set *M*_lr_ = 900, with 30 uniform samples in *µ*FA and D_iso_, respectively, and use *M*_**u**_ = 768 samples on the unit sphere based on the equal-area HEALPix Gorski et al. (2005) method. Based on the estimation results, we compute the FOD using Eq. (13) and select the *K* peaks and their corresponding peak orientations.

The second step refines the microstructure estimates using a high-resolution grid with *M*_hr_ samples in the *µ*FA − D_iso_ plane and the fixed *K* fiber-orientation samples, giving *M*_2_ = *M*_hr_*K* total samples. In this work, we set *M*_hr_ = 40000, with 200 samples in *µ*FA and D_iso_, respectively. We note that this assumption is feasible not only for modeling WM fiber bundles to investigate fiber-specific microstructure, but also for modeling GM microstructure with near-isotropic components using a suitable *P* (*µ*FA, D_iso_ | **u**). Based on this assumption, we introduce the following adaptive sampling and estimation method to recursively estimate fiber orientation and microstructure.

## 3. Experiments

The performance of the MaxEnt-DTD method was evaluated using in vivo and synthetic data with standard linear diffusion encoding, as used in the Human Connectome Project (HCP) (Sotiropoulos et al., 2013), and B-tensor diffusion encoding methods (Westin et al., 2016; Szczepankiewicz et al., 2019).

### 3.1. In vivo data

#### Human Connectome Project (HCP)

The HCP diffusion MRI data (Van Essen et al., 2013; Sotiropoulos et al., 2013) were acquired on a customized Siemens 3T scanner equipped with 100 mT/m gradients using iPAT=2, multiband factor=3, TR/TE=5520/89.5 ms, and 1.25 mm isotropic voxels. Diffusion-weighted images were acquired with *b* = 1000, 2000, and 3000 s/mm^2^, 90 gradient directions per shell, and 18 non-diffusion-weighted volumes. In this dataset, the B-tensors were of rank 1 (linear tensor encoding, LTE). FreeSurfer (Fischl, 2012) tissue segmentations were derived from T1-weighted MRI and co-registered to dMRI to obtain tissue label maps for region-based analysis.

#### Tensor-valued encoding (B-tensor) data

The proposed method was also applied to a public tensor-valued dMRI dataset provided by (Szczepankiewicz et al., 2019), acquired in a healthy human brain on a 3T MAGNETOM Prisma system (Siemens Healthcare). Data were acquired using numerically optimized, Maxwell-compensated free gradient waveforms at 2.5 mm isotropic resolution, with an acquisition matrix of 92 × 92 × 25, TR/TE = 3200/92 ms, and iPAT=2.

The diffusion encoding protocol comprised linear (LTE), planar (PTE), and spherical (STE) B-tensor waveforms at four diffusion-weighting levels, *b* = 100, 700, 1400, and 2000 s/mm^2^. LTE and PTE were each sampled with 10, 10, 16, and 46 rotations across the four b-values (82 volumes per encoding shape), whereas STE was sampled with 10 rotations per b-value, each repeated five times (200 volumes). Together with 13 non-diffusion-weighted images, this yielded a total of 377 volumes per voxel. The provided data were processed with motion and eddy-current correction.

### 3.2. Synthetic data

To enable quantitative benchmarking, we generated synthetic datasets using a modified standard model (Novikov et al., 2018) following the setup of (Consagra et al., 2025). The synthetic model consisted of one to three fiber bundles, where each fiber population was assigned a ground-truth (*µ*FA, D_iso_) pair drawn from physiologically plausible ranges (*µ*FA ∈ [0.3, 0.9], D_iso_ ∈ [0.5, 1.5] × 10^−3^ mm^2^*/*s) and an orientation specified by a unit vector. In cases with more than one fiber bundle, the crossing angle ranged from 10^◦^ to 90^◦^. The signal from each population was generated using the axisymmetric kernel of Eq. (5), and the population signals were combined according to prescribed volume fractions summing to one. The ground-truth DTD was therefore represented as a weighted sum of delta functions in the (*µ*FA, D_iso_, *ϕ, ψ*) space.

Signals were synthesized for two acquisition schemes consistent with the in vivo analyses: (i) standard multi-shell LTE (*b* = 1000, 2000, 3000 s/mm^2^, 90 directions per shell), and (ii) tensor-valued encoding (LTE, PTE, and STE up to *b* = 2000 s/mm^2^). The synthetic signals were then corrupted with additive Gaussian noise whose variance was calibrated to yield target SNR levels of 10, 20, 50, and 100 at the highest b-value. A total of 25,500 test samples were generated for each protocol for quantitative evaluation.

### 3.3. Evaluation metrics

The performance of MaxEnt-DTD was evaluated in three aspects using fiber orientation distribution, fiber-specific *µ*FA −D_iso_ distribution and the rotation invariant microstructure metrics derived from the cumulants of the DTD. For synthetic experiments, where ground truth is available, all metrics are reported quantitatively at the voxel level. For in vivo experiments, we use region-of-interest (ROI) analyses and qualitatively evaluate the microstructure measures.

For comparison, we applied several versions of the CSD approach (Tournier et al., 2007; Descoteaux et al., 2008; Jeurissen et al., 2014, 2019; Raffelt et al., 2017), generalized CSD with B-tensor encoding waveforms (Jeurissen and Szczepankiewicz, 2021; Karan et al., 2022) for FOD estimation. We examined the trade-off in performance by using different subsets of the dMRI samples for both the HCP-type and B-tensor encoding protocols. For consistent comparison, the FOD for MaxEnd-DTD were filterd using 8th order spherical harmonics as used by the CSD methods.

We compared the performance of the MaxEnt-DTD with results from the Monte-Carlo method (Topgaard, 2019). We applied the same algorithm as proposed in (Topgaard, 2019) with the same signal presentation as the MaxEnt-DTD methods.

We also examined the QTI rotation invariant microstructure measures proposed in (Westin et al., 2016) based on the estimated cumulants of MaxEnt-DTD. We compared the performance with the results from the MD-dMRI toolbox (Nilsson et al., 2018). Below are the specific evaluation metrics. In theory, the HCP protocl does not provide information to estimate the second-order cumulant which involves 21 parameters using the method in (Westin et al., 2016) while the HCP protocol only provides 15 degree of freedoms. However, axisymetric parmetriziation of the microstructural tensors reduces the number of parameters for the second order cumulants to 10 which may improve the reliability for DTD-based sampling methods. The comparison provides insights to understand the the performances these methods with varying protocols.

#### 3.3.1. Fiber orientation evaluation

##### Angular error (AE)

We use the same metrics proposed in (Consagra et al., 2025) to evaluate the accuracy of FOD estimation. For a voxel with *N*_gt_ ground-truth fiber orientations 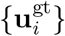 and *N*_est_ estimated peaks 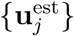 extracted from the marginal FOD, the angular error is computed as the mean of the minimum angular deviation between matched peak pairs:

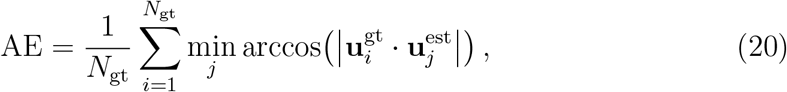

where the absolute value accounts for the antipodal symmetry of diffusion tensor orientations. Lower AE indicates more accurate orientation recovery.

##### Percentage of correct peaks (PCP)

To evaluate the reliability of peak detection in the presence of spurious or missed peaks, we report the percentage of correct peaks (PCP), defined as the proportion of voxels in which all *N*_gt_ ground-truth peaks are recovered within an angular tolerance of *τ*_ang_ = 20^◦^, and no spurious peaks are detected. Higher PCP indicates more robust peak detection. The tolerance is chosen to be smaller than the smallest inter-fiber angle evaluated in our experiments, ensuring that PCP captures genuine identification accuracy.

#### 3.3.2. Fiber-specific microstructure evaluation

##### Wasserstein distance (WD)

To quantify the discrepancy between an estimated DTD 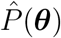 and the ground-truth distribution *P* ^gt^(***θ***), we use the *p*-Wasserstein distance (Villani, 2009; Peyré et al., 2019):

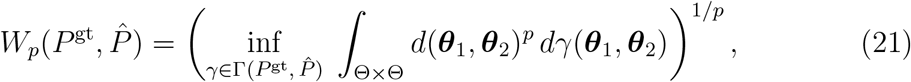

where 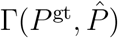 is the set of all joint distributions on Θ × Θ with marginals *P* ^gt^ and 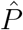, and *d*(***θ***_1_, ***θ***_2_) is a ground distance combining a Euclidean component in the spectral domain (*µ*FA, *D*_iso_) with a geodesic component on the orientation domain **u** ∈ S^2^. We use *p* = 1 on the marginal distributions *P* (**u**) and *P* (*µ*FA, *D*_iso_), which is equivalent to the Earth Mover’s distance(Rubner et al., 2000), and *p* = 2 on the full four-dimensional distribution, which more heavily penalizes outlier mass placements. Lower WD indicates better distributional agreement with the ground truth.

#### 3.3.3. Rotation-invariant microstructure measures

##### Pearson correlation (ρ) and concordance correlation coefficient (CCC)

To assess the recovery of voxel-level scalar metrics derived from the estimated DTD, including the MD, FA, microscopic FA (*µ*FA), mean kurtosis (MK), and the isotropic and anisotropic components of microscopic kurtosis (MK_*i*_ and MK_*ad*_), against their ground-truth counterparts, we compute two complementary measures on synthetic data generated using both the HCP multi-shell acquisition and the B-tensor protocol. Given paired ground-truth and predicted values 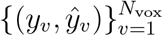, the Pearson correlation coefficient

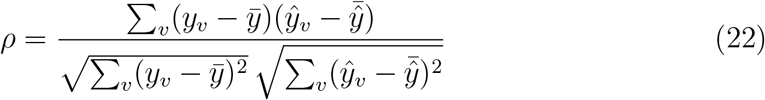

measures the precision of the linear association, independent of global scale and offset. The concordance correlation coefficient (Lawrence and Lin, 1989)

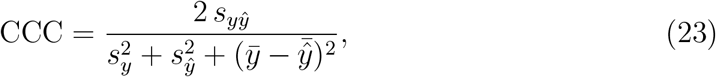

where *s*_*yŷ*_ is the sample covariance and 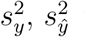 the sample variances, additionally quantifies agreement with the identity line *ŷ* = *y* by penalizing systematic bias and scaling errors. CCC factorizes as CCC = *ρ C*_*b*_, where the bias-correction factor *C*_*b*_ ∈ (0, 1] measures how far the best-fit line departs from the identity line; reporting *ρ* and CCC together therefore disentangles *precision* (ranking fidelity) from *accuracy* (absolute calibration). Both coefficients lie in [−1, 1], with values approaching unity indicating better recovery. To characterize robustness across acquisition conditions, we report *ρ* and CCC as a function of signal-to-noise ratio (SNR) over levels ranging from 10 to 100, using simulated data based on both the HCP and B-tensor protocols.

### 4. Results

#### 4.1. Results on synthetic data

#### 4.1.1. Fiber orientation estimation

Fig. 1 reports the angular error (AE) and proportion of correct peaks (PCP) on synthetic data generated under the HCP protocol. Fig. 1A shows that MaxEnt-DTD with all multi-shell data achieved the lowest AE. MaxEnt-DTD also achieved much lower AE than CSD even when only single-shell data were used. Fig. 1D shows that MaxEnt-DTD with multi-shell data achieved the highest PCP, with the CSD methods producing similar results. For single-shell data with *b* = 1000 s/mm^2^, CSD achieved a PCP of approximately 0.70, compared with 0.90 for MaxEnt-DTD. For higher b-values or multi-shell data, the PCP values for CSD and MaxEnt-DTD were all greater than 0.90, whereas the PCP for MC was much lower, at approximately 0.60.

**Figure 1:**
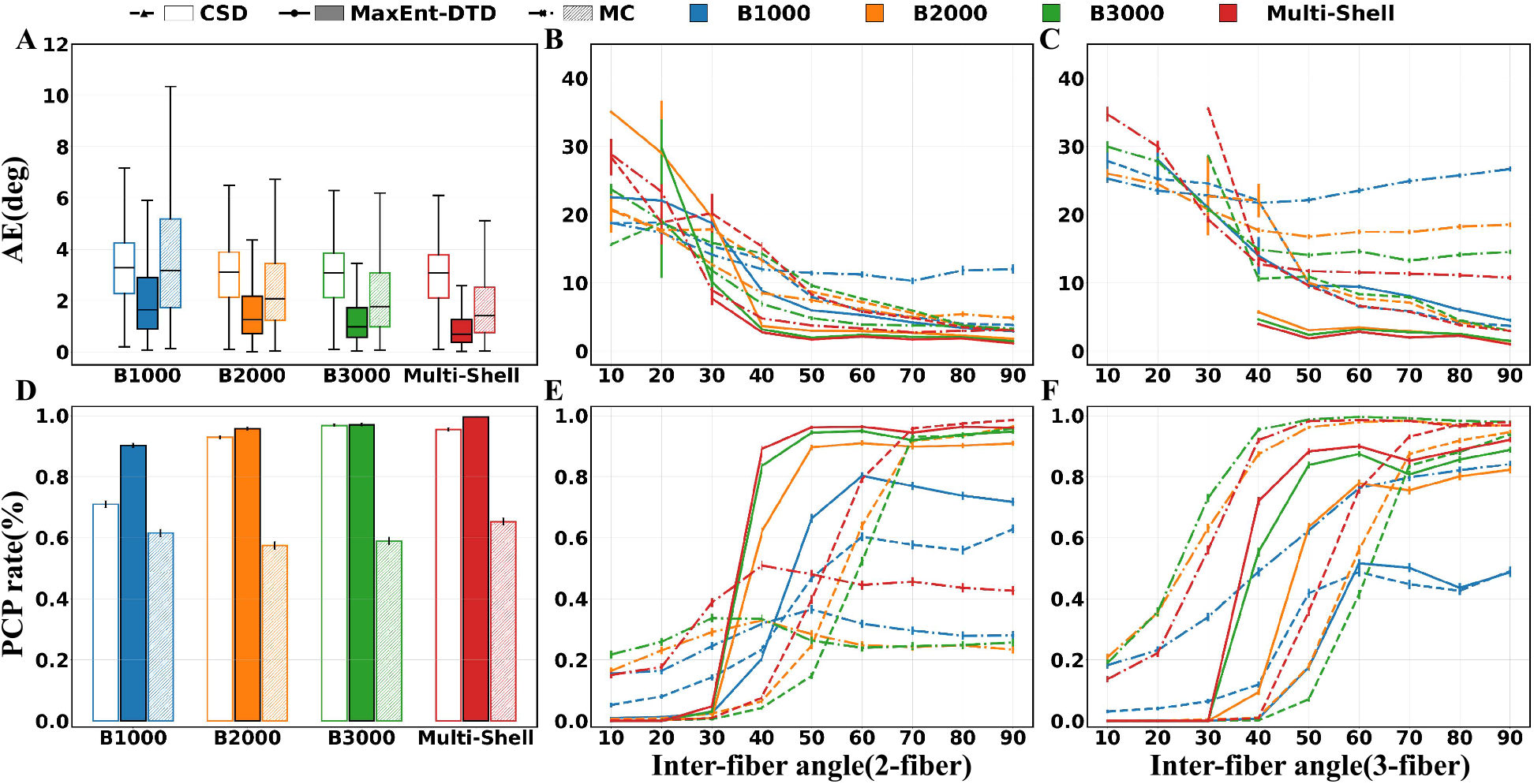
(Synthetic data with HCP protocol) A: AE for 1-fiber configurations shown as boxplots grouped by single-shell and Multi-shell conditions for the CSD and Proposed methods; B: Mean AE for 2-fiber; C: 3-fiber configurations plotted against inter-fiber angles ranging from 10° to 90°. D: PCP rate for 1-fiber configurations displayed as grouped bar plots; E: Mean PCP rate for 2-fiber; F: 3-fiber configurations evaluated across varying inter-fiber angles. Error bars represent the standard error of the mean (SEM) across samples within each angle bin for AE curves, and the binomial SEM for PCP plots.

The second and third columns of Fig. 1 illustrate the AE and PCP for two- and three-fiber cases with varying crossing angles. MaxEnt-DTD consistently achieved lower AE and higher PCP than CSD across all encoding settings and inter-fiber angles. The angular resolution limit, defined as the smallest inter-fiber angle at which the correct number of peaks was reliably detected, improved from approximately 40-50^◦^ for CSD to approximately 30^◦^ for MaxEnt-DTD using multi-shell data, indicating an approximately 10-20^◦^ improvement in angular resolution. Similar results were observed for three-fiber cases, although the PCP for MaxEnt-DTD was slightly lower, decreasing from approximately 0.9 to 0.7 at 40^◦^. Although MC showed higher PCP in Fig. 1F, it was associated with high AE, as shown in Fig. 1C, indicating that MC likely detected a large number of false-positive peaks.

Fig. 2 shows the angular performance of the methods for synthetic data generated under the B-tensor encoding protocol. Overall, the results were similar to those shown in Fig. 1. The first column shows the single-fiber results for different subsets of samples, including LTE-only, PTE-only, and the full dataset. Overall, MaxEnt-DTD with the full dataset achieved the most accurate angular performance. The second and third columns show the results for two- and three-fiber cases. The comparison between Fig. 2(E,F) and Fig. 1(E,F) shows that the HCP protocol provided better angular performance than the B-tensor encoding protocol. Specifically, the B-tensor encoding data yielded PCP values of 0.95 and 0.72 for two- and three-fiber cases with 50^◦^ crossing angles, whereas the corresponding PCP values for the HCP protocol were 0.96 and 0.88, respectively. The MC method still showed a high false-positive rate, characterized by both high PCP and high AE.

**Figure 2:**
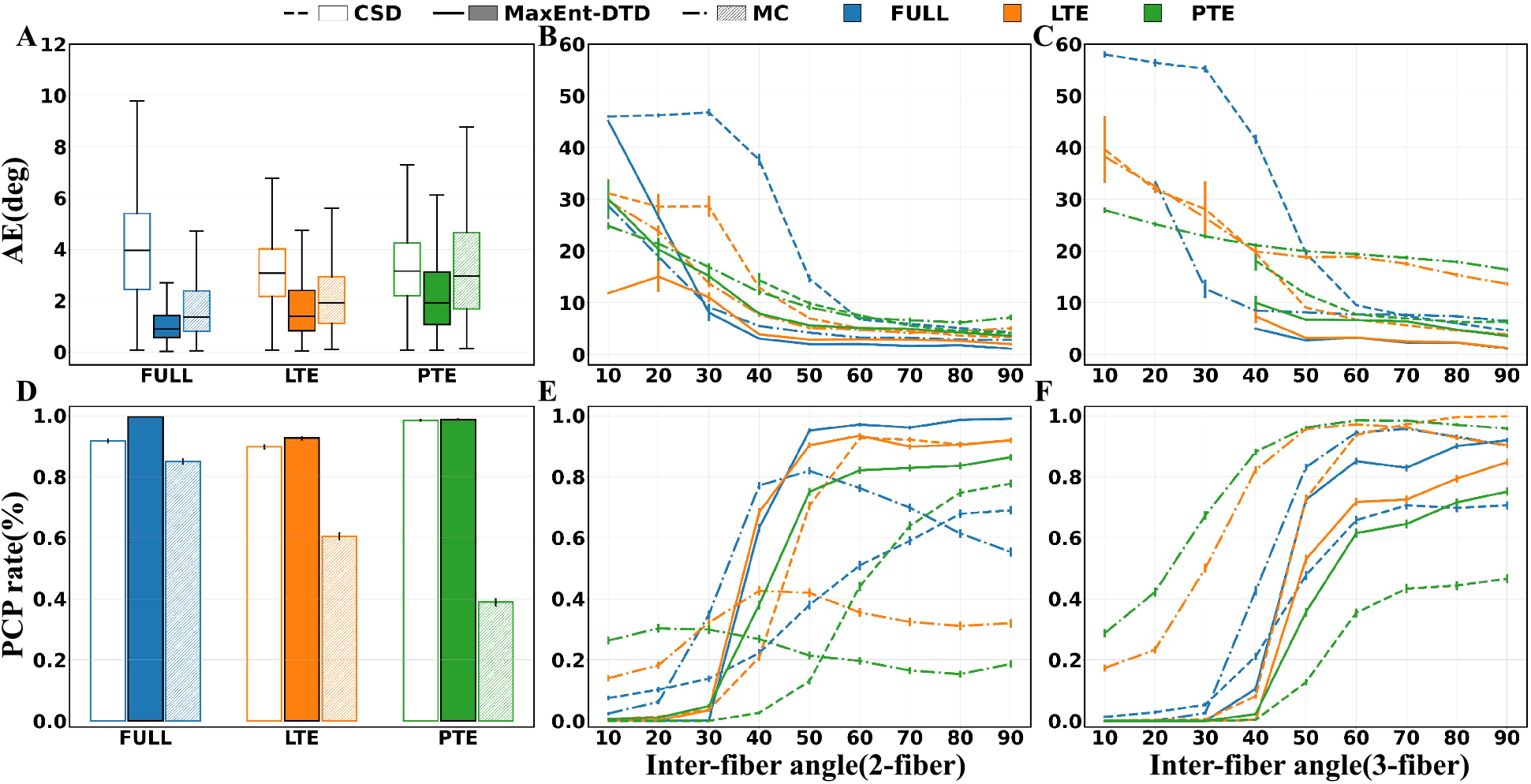
(Synthetic data with B-tensor encoding) A: AE for 1-fiber configurations shown as box-plots grouped by FULL, LTE, and PTE encodings for CSD and Proposed methods; B: Mean AE for 2-fiber; C: 3-fiber configurations plotted against inter-fiber angles from 10° to 90°; D: PCP for 1-fiber configurations displayed as bar plots; E: Mean PCP rate for 2-fiber; F: 3-fiber configurations evaluated across varying inter-fiber angles.

#### 4.1.2. Fiber-specific microstructure estimation

Fig.3 illustrates the Wasserstein distance, i.e., EMD, between the estimated DTD and the ground-truth distribution, evaluated for single-, two-, and three-fiber configurations under four encoding settings (multi-shell, B1000, B2000, B3000). Lower values indicate closer agreement with the ground truth and better performance.

**Figure 3:**
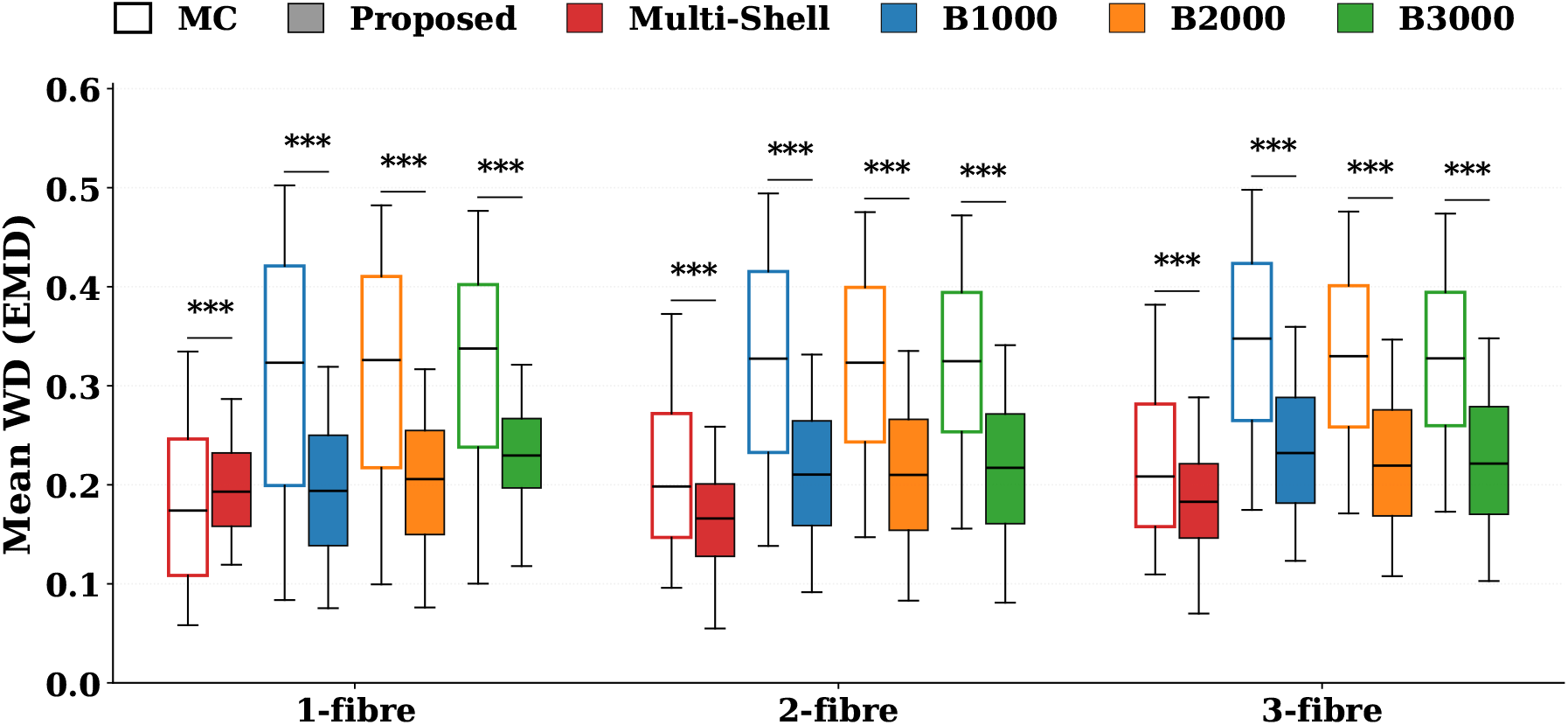
Microstructure estimation accuracy on synthetic data with HCP protocol, quantified by the Wasserstein distance (WD) between the estimated DTD and the ground truth (lower is better). Comparison of Monte Carlo (MC, open) and MaxEnt-DTD (filled) for single-, two-, and three-fiber configurations under multi-shell and single-shell encodings. The notation “***” denotes *p <* 0.001 (Wilcoxon signed-rank test).

For the single-fiber case (Fig.3, left), MaxEnt-DTD and Monte Carlo (MC) inversion achieve comparable accuracy under multi-shell encoding (median WD of 0.20 versus 0.17). Under each single-shell setting, however, MaxEnt-DTD substantially outperforms MC, with median WD reduced from approximately 0.32 to 0.20 at *b* = 1000 s/mm^2^ and similar improvements at *b* = 2000 and *b* = 3000 s/mm^2^ (*p <* 0.001, Wilcoxon signed-rank test).

For the two- and three-fiber cases (Fig.3, middle and right), MaxEnt-DTD consistently achieves lower median WD than MC across all four encoding settings, including multi-shell. The MC distributions exhibit substantially wider interquartile ranges, indicating reduced reliability when encoding diversity is limited. The gap between the two methods narrows but remains significant under multi-shell encoding (*p <* 0.001 for both two- and three-fiber configurations). Across all fiber multiplicities, MaxEnt-DTD with multi-shell encoding yields the lowest median WD, demonstrating that the method benefits from richer encoding while remaining accurate under reduced acquisition protocols.

Fig.4 shows the corresponding Wasserstein-distance comparison on synthetic tensor-valued encoding data under FULL, LTE, and PTE acquisitions. Overall, the MaxEnt and MC methods have similar performance when FULL samples are used. The variance of the MC methods increases dramatically with fewer samples. Moreover, the performance of both methods is much worse for multi-fiber cases than results for single-fiber cases.

**Figure 4:**
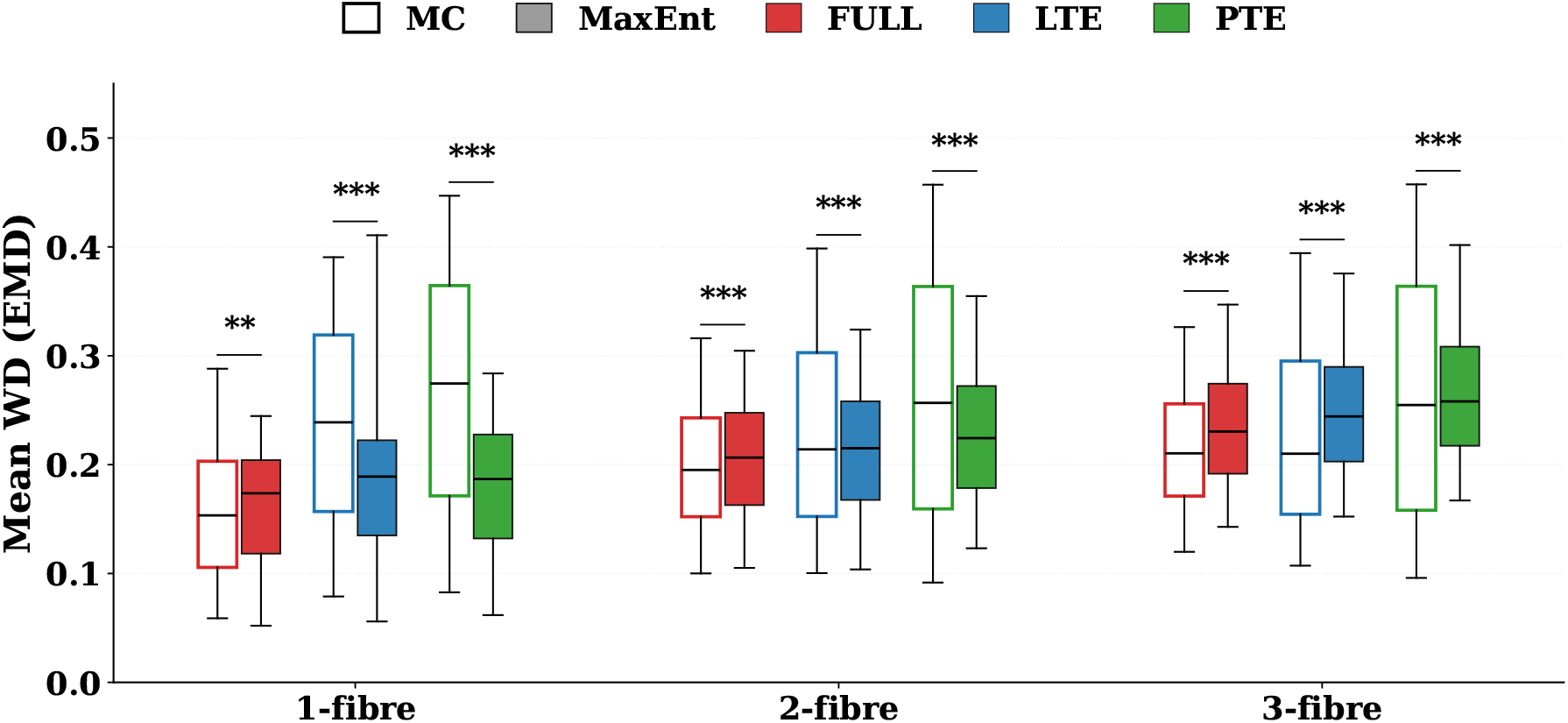
Microstructure estimation accuracy on synthetic data with B-tensor encoding, quantified by the Wasserstein distance (WD) between the estimated DTD and the ground truth (lower is better). Comparison of Monte Carlo (MC) and MaxEnt-DTD under FULL, LTE, and PTE encoding settings.

#### 4.1.3. Rotation-invariant microstructure measures

Figs.5 and 6 illustrate the estimation accuracy for six rotation-invariant scalar metrics, including MD, FA, *µ*FA, MK, MK_I_, and MK_A_, computed from the DTD cumulants using the HCP and B-tensor encoding protocols, respectively. For MaxEnt-DTD and MC methods, the cumulants were computed by numerical sampling based on Eq. (15). In contrast, the MD-dMRI method directly computed the cumulants without estimating the DTD based on (Westin et al., 2016). The two figures show results for the HCP linear-encoding protocol and the B-tensor encoding protocol, respectively. The top panel (A) of each figure, including the first two rows, shows redicted-versus-ground-truth scatter with the Pearson correlation (*ρ*) and the concordance correlation coefficient (CCC) annotated for synthetic data with SNR=30. The bottom rows (B) report *ρ* and CCC as functions of SNR from 10 to 100.

**Figure 5:**
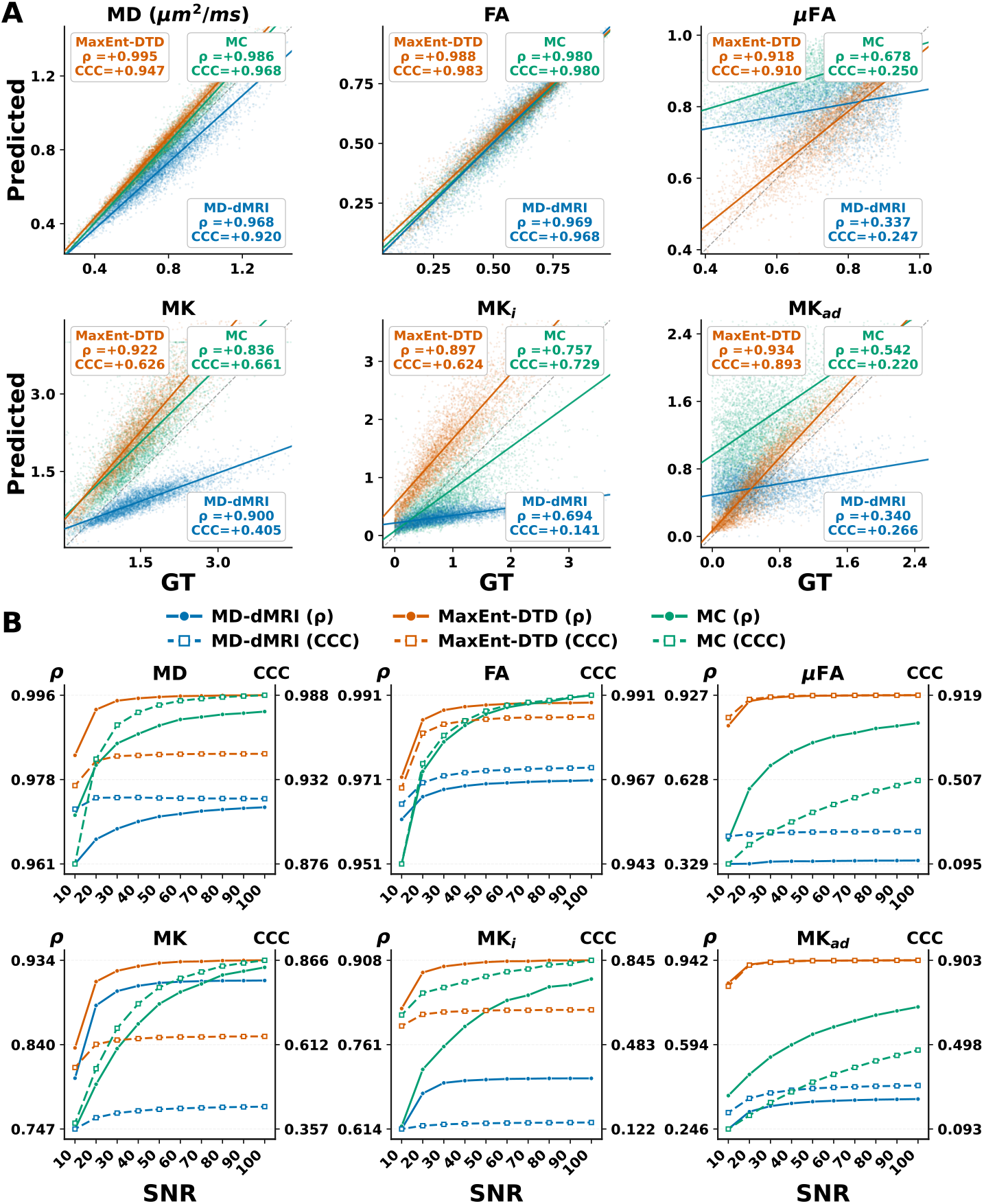
Performance on rotation-invariant microstructure metrics using HCP protocol: (A) Predicted-versus-ground-truth scatter at SNR = 30 for MD, FA, *µ*FA, MK, MK_*i*_, and MK_*ad*_, comparing MD-dMRI (blue), MaxEnt-DTD (orange), and MC (green); the dashed gray line is the identity, solid lines are least-squares fits, and the annotated *ρ* and CCC are the Pearson and concordance correlation coefficients. (B) *ρ* (solid, left axis) and CCC (dashed, right axis) as functions of SNR from 10 to 100 for each metric.

**Figure 6:**
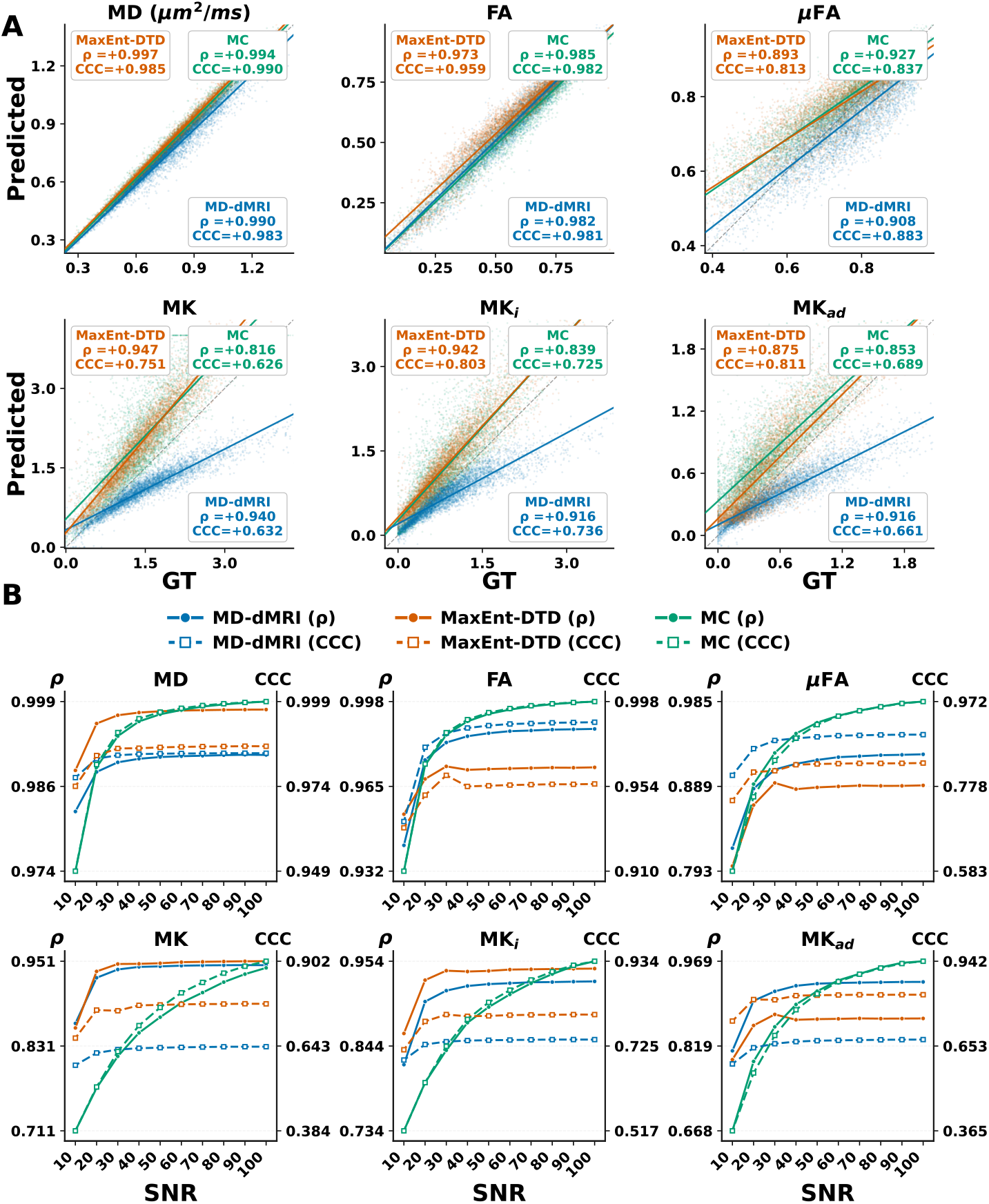
Performance on rotation-invariant microstructure metrics using B-tensor encoding: Panel (A) shows the predicted-versus-ground-truth scatter at SNR = 30; panels and conventions otherwise as in Fig.5.

For the HCP-protocol results (Fig.5), all three methods accurately recovered MD and FA. The MaxEnt and MC methods have similar values for *ρ* and CCC. MD-dMRI produced similar but slightly lower correlations, with *ρ* = 0.968 and 0.969 and CCC = 0.920 and 0.968 for MD and FA, respectively. The estimates were robust across SNR levels, but MaxEnt-DTD and MC generally had slightly higher *ρ* and CCC values than MD-dMRI, as shown in the lower panels.

The MaxEnt-DTD preserved a stronger correlation with the *µ*FA measures than the other two methods, with *ρ* = 0.918, CCC = 0.910. The biased results by the MD-dMRI were expected because the HCP protocol does not provide sufficient information to measure all second-order cumulants (Westin et al., 2016). The improved performance of MaxEnt-DTD is related to the simplified axisymmetric structure of the microscopic diffusion tensors and the angular resolution provided by DTD estimation. On the other hand, although MC also imposed an axisymmetric structure, its reduced performance was likely due to its increased sensitivity to noise and biased estimation of fiber orientations.

The MK and MK_I_ results showed similar patterns: MaxEnt-DTD achieved the highest *ρ* and CCC values and remained robust at lower SNR, whereas MC performance decreased substantially at low SNR. MaxEnt-DTD slightly overestimated high MK and MK_I_ values, whereas MD-dMRI significantly underestimated the MK and MK_I_ metrics. The increased bias and randomness in the estimation results for the MC and MD-dMRI methods are related to the limited informaiton in the HCP protocol.information in the HCP protocol.

Fig.6 shows the B-tensor encoding results. Overall, the B-tensor encoding method significantly improved the prediction of *µ*FA and MK_A_ measures, especially for the MD-dMRI and MC methods. MaxEnt-DTD showed better performance metrics than the other two methods, especially at relatively low SNR values between 30 and 50. The main differences from the HCP-protocol results in Fig.5 appeared in the *µ*FA and MK_A_ plots. All three methods performed substantially better in estimating *µ*FA because of the additional information probed by B-tensor encodings. MD-dMRI showed the most accurate estimation for MK_A_, whereas the other two methods had similar performance.

The lower panels of the Figs.5 and 6 show the dependence of the performance metrics of the three methods on the SNR to demonstrate their sensitivity to noise. The MC method showed strong dependence on measurement noise, with significantly lower performance than the other two methods when the SNR is lower than 30. While the other two methods showed more robust performance at relatively lower SNR. The reliable performance of the MD-dMRI method is due to its characterization of finite cumulants. The improved robustness of the MaxEnt-DTD is related to the explicit incorporation of measurement noise in the estimation algorithm.

### 4.2. Results on HCP data

#### 4.2.1. Fiber orientation estimation

Fig.7 compares the FOD estimated from HCP data by the proposed method with CSD reconstructions under three shell settings: single-shell CSD at *b* = 1000 s/mm^2^ (B1000), single-shell CSD at *b* = 3000 s/mm^2^ (B3000), and multi-shell CSD (B1000, B2000, B3000). The whole-brain FOD field in Fig.7A shows that the proposed method produces spatially coherent orientation patterns broadly consistent with the Multi-shell result, whereas both single-shell reconstructions exhibit reduced orientational detail, particularly in regions of complex fiber configurations.

**Figure 7:**
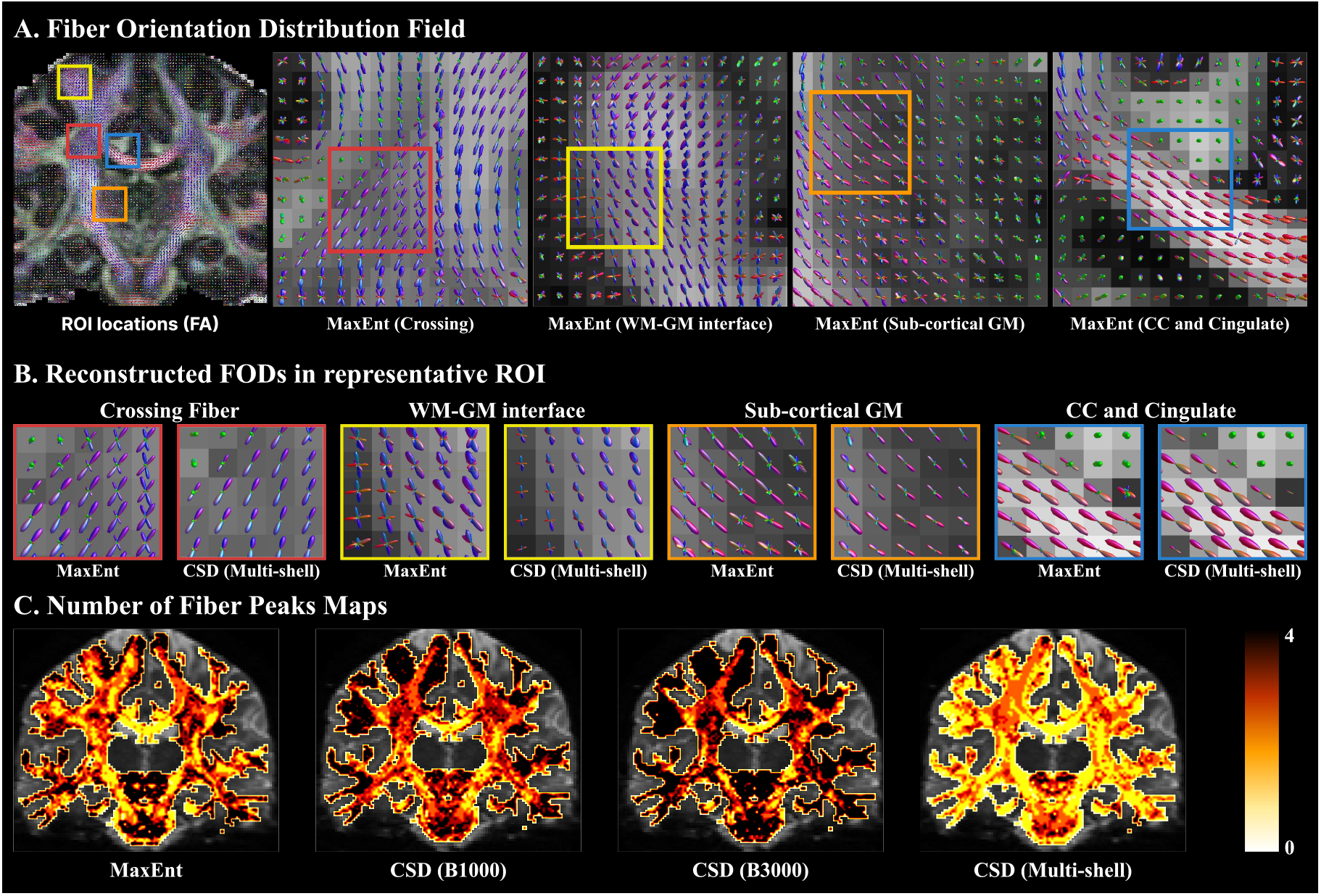
Comparison of FOD reconstructions and estimated fiber peaks across different acquisition settings. (A) FOD fields generated using the Proposed, B1000, B3000, and Multi-shell methods, overlaid on an anatomical background. Colored boxes outline selected ROIs. (B) Magnified views of the reconstructed FOD within the representative ROIs: Crossing Fiber (red), WM-GM interface (yellow), sub-cortical GM (orange), and CC and Cingulate (blue), comparing the four respective settings. (C) Heat maps illustrating the estimated number of fiber peaks for each corresponding method, with the color scale ranging from 0 to 4.

Magnified views of FODs in four representative regions of interest (ROIs) are shown in Fig.7B. In the crossing-fiber ROI, the proposed method recovers FODs with two to three well-separated orientations in a voxel. In contrast, the multi-shell CSD reconstruction tends to produce smoother, less angularly resolved configurations. At the WM and GM interface, the proposed method produces FODs that transition from elongated lobes in deep white matter to compact profiles near the cortical surface, whereas the other reconstruction yields barely visible FOD glyphs in this region. In sub-cortical gray matter, the proposed method generates small, low-amplitude FODs. The multi-shell CSD yields comparable low-amplitude patterns. In the corpus callosum and cingulate region, both methods recover predominantly unimodal orientations that are broadly consistent with known anatomy, while the proposed method yields visually sharper lobe profiles.

Fig.7C shows the maps for the estimated number of fiber orientations for the MaxEnt method, CSD with single-shell data for three b-values and multi-shell data. The proposed method identifies a higher density of multi-peak voxels throughout white matter relative to both single-shell settings, highlights the junction of the corticospinal tract (CST), superior longitudinal fasciculus (SLF), and corpus callosum (CC), where the anatomical region reveals a three-bundle crossing fiber configuration, and remains in close agreement with the Multi-shell CSD map.

#### 4.2.2. Fiber-specific microstructure estimation

Fig. 8 shows the estimated fiber-specific *P* (*µ*FA, D_iso_ | **u**) conditioned on peak orientations in representative brain regions. Overall, each white-matter fiber bundle is associated with two dominant components: one with high *µ*FA and low D_iso_, and another with lower *µ*FA and higher D_iso_, as shown in Fig. 8(A,C,E). Gray-matter regions are associated with low *µ*FA and high D_iso_, as shown in Fig. 8(B), but they also include broad distributions that extend into high-*µ*FA regions. The CSF region includes only one dominant distribution around the highest D_iso_ value with *µ*FA = 0, as shown in Fig. 8(D), as expected.

**Figure 8:**
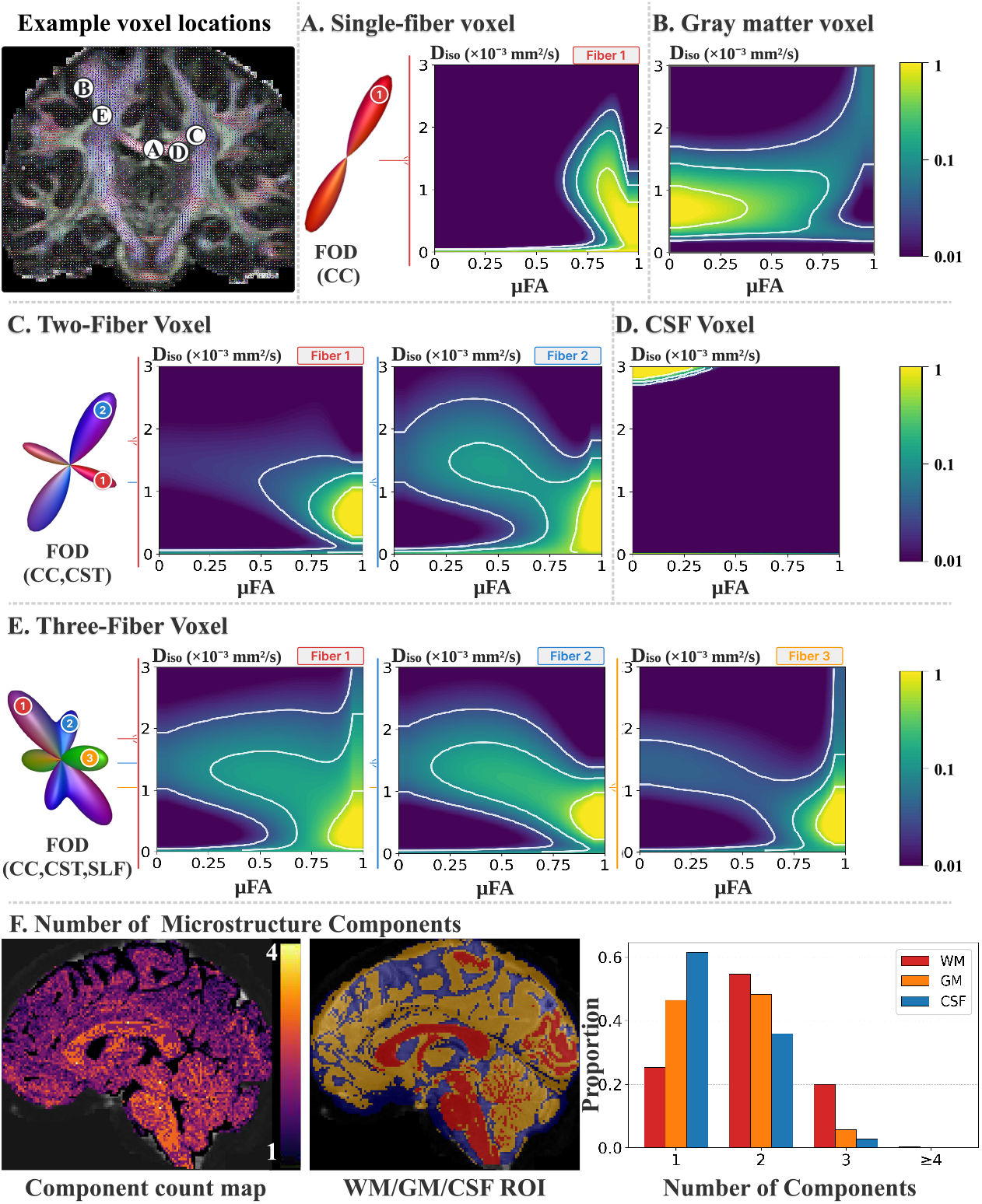
Fiber-specific microstructure by MaxEnt-DTD in representative in vivo voxels (HCP). *Top left:* five voxels (A-E) marked on the FOD field (positions approximate). Each panel shows the FOD and the recovered distribution over *µ*FA and D_iso_; log-scaled density (a.u.) with white contours. For WM voxels (A, C, E) the per-fiber *P* (*µ*FA, D_iso_ | **u**) is shown (Fiber 1, red; 2, blue; 3, orange); for GM (B) and CSF (D) the voxel-level *P* (*µ*FA, D_iso_). (A) Single-fiber (CC); (B) gray matter; (C) two-fiber (CC, CST); (D) CSF; (E) three-fiber (CC, CST, SLF). (F)(Left) The average of components, i.e., the number of peaks of *µ*FA and D_iso_, over all peak orientation directions. (Right) The histogram of the number of components over different tissue regions for the tissue segmentation map shown in the middle panel.

The last row of Fig. 8 shows the number of components, i.e. the number of peaks in *P* (*µ*FA, D_iso_ | **u**) averaged over the peak orientations. The left panel shows that most single-fiber white-matter voxels include three components, whereas other areas include fewer components. The right panel shows the histogram of the number of components in different brain-tissue regions based on the tissue label maps shown in the center image. In general, most white-matter areas include two components, whereas gray-matter areas have one or two components.

#### 4.2.3. Cumulant-based rotation-invariant metrics

Fig.9 shows maps of the estimated rotation-invariant microstructure measures. The three rows correspond to MaxEnt-DTD, MD-dMRI, and MC, respectively. The three methods produced similar FA and MD maps. The MC method produced much higher values for *µ*FA than the other two methods across the whole brain. The MD-dMRI method also showed elevated *µ*FA characterized by high *µ*FA in CSF areas which is consistent to the overestimation results in synthetic data related to the limited information of the HCP protocol. The MaxEnt method showed similar constrast in *µ*FA as the MD-dMRI but with more accurate values in the CSF areas and overall noisier maps.

**Figure 9:**
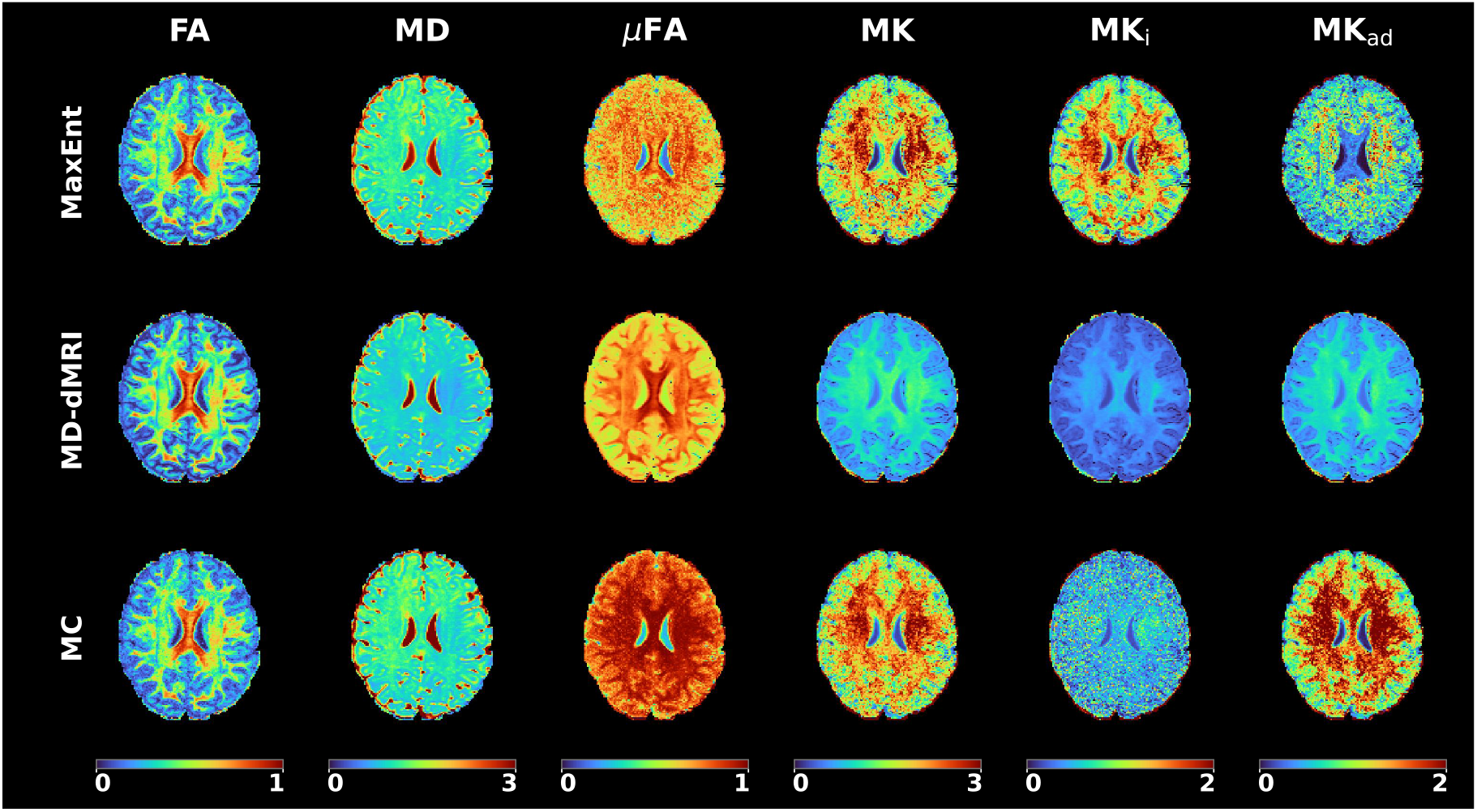
Rotation-invariant microstructure measures (HCP). Rows: MaxEnt-DTD (Max-Ent), MD-dMRI, MC. Columns: fractional anisotropy (FA), mean diffusivity (MD, ×10^−3^ mm^2^*/*s), microscopic FA (*µ*FA), mean kurtosis (MK), and its isotropic and anisotropic parts (MK_i_, MK_ad_).

The MD-dMRI method showed significantly lower MK metrics than the other two methods, consistent to results in Fig. 1. Overall, these trends are consistent with Fig.5, where MaxEnt-DTD tended to overestimate high MK values and MD-dMRI tended to underestimate MK. The MC-based MK maps also appeared noisier, especially in the MK_I_ map, than those from the other methods, likely reflecting greater sensitivity to measurement noise.

### 4.3. Results on in vivo B-tensor dMRI

#### 4.3.1. Fiber orientation estimation

Fig.10 compares the FODs estimated from tensor-valued diffusion encoding data by the proposed method and three encoding shape specific CSD variants: Full-CSD (joint LTE/PTE/STE), LTE-CSD (linear tensor encoding only), and PTE-CSD (pla-nar tensor encoding only). In the whole-brain FOD field in Fig.10A, the proposed method yields spatially coherent orientation patterns with sharper lobe profiles than the CSD reconstructions, most notably in regions containing multiple crossing fiber populations.

**Figure 10:**
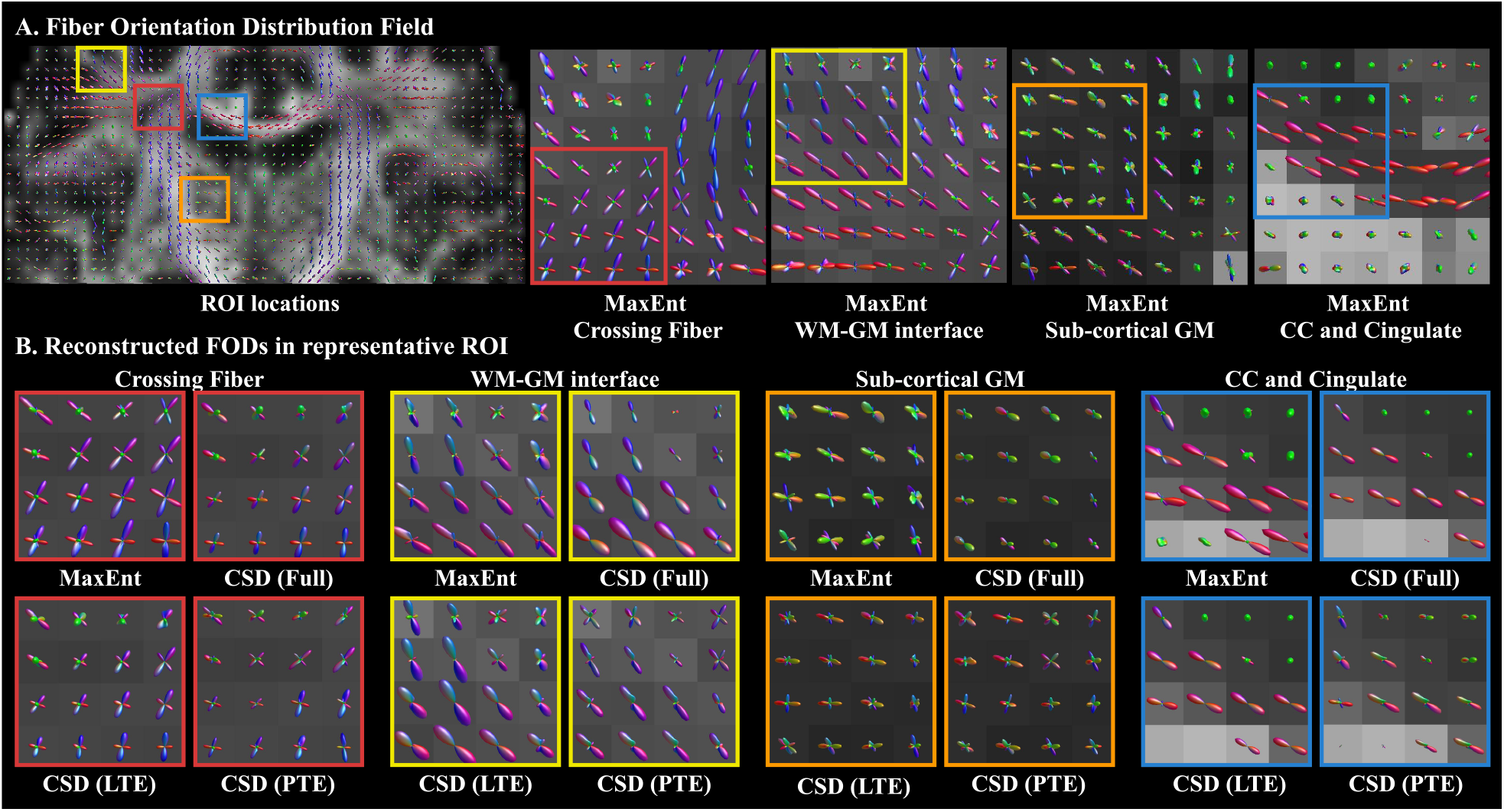
Comparison of FOD reconstruction methods. (A) FOD fields generated by the Proposed method, Full-CSD, LTE-CSD, and PTE-CSD overlaid on an anatomical background. Colored boxes indicate ROIs selected for detailed inspection. (B) Magnified views of reconstructed FODs in four distinct anatomical regions: crossing fibers (red), WM-GM interface (yellow), sub-cortical GM (orange), and the CC and cingulate (blue). Each panel provides a visual comparison of the angular resolution and noise profiles of the Proposed method against the Full-CSD, LTE-CSD, and PTE-CSD frameworks.

The magnified views of FODs for four representative ROIs are shown in Fig.10B. In the crossing-fiber ROI, the proposed method produces well-defined multi-peak FODs, while Full-CSD yields comparable directional structure with broader lobes. The single-encoding variants (LTE-CSD and PTE-CSD) appear less angularly resolved in this region, suggesting that the combination of multiple encoding shapes contributes to improved angular discrimination. At the WM-GM interface, the proposed method resolves both deep white matter fibers and superficial white matter fiber crossings, with FODs transitioning from multi-peak profiles in the deep white matter to compact configurations near the cortical surface. In contrast, Full-CSD yields inflated FOD glyphs with spurious lobes in this region, while LTE-CSD and PTE-CSD produce attenuated FODs that do not clearly delineate the superficial fiber crossings. In sub-cortical gray matter, all methods yield low-amplitude, approximately isotropic FODs, though the CSD variants occasionally exhibit residual directional structure. In the corpus callosum and cingulate region, all four methods recover unimodal callosal orientations consistent with known anatomy. The proposed method and Full-CSD additionally resolve the cingulate fiber population, which appears attenuated in the LTE-CSD and PTE-CSD results.

#### 4.3.2. Fiber-specific microstructure estimation

Fig.11 illustrates the fiber-specific microstructure measures for the B-tensor encoding dataset. Fig.11(A,C,E) shows the *µ*FA-D_iso_ distributions conditioned on the fiber orientations for single-fiber, two-fiber, and three-fiber voxels, respectively. Similar to the results in Fig.8, these distributions included a dominant component with high *µ*FA and low D_iso_. But a major difference is a component with low *µ*FA around zeros that are not shown in results in 8 which may be related to new microstructure information probed by B-tensor encodings.

**Figure 11:**
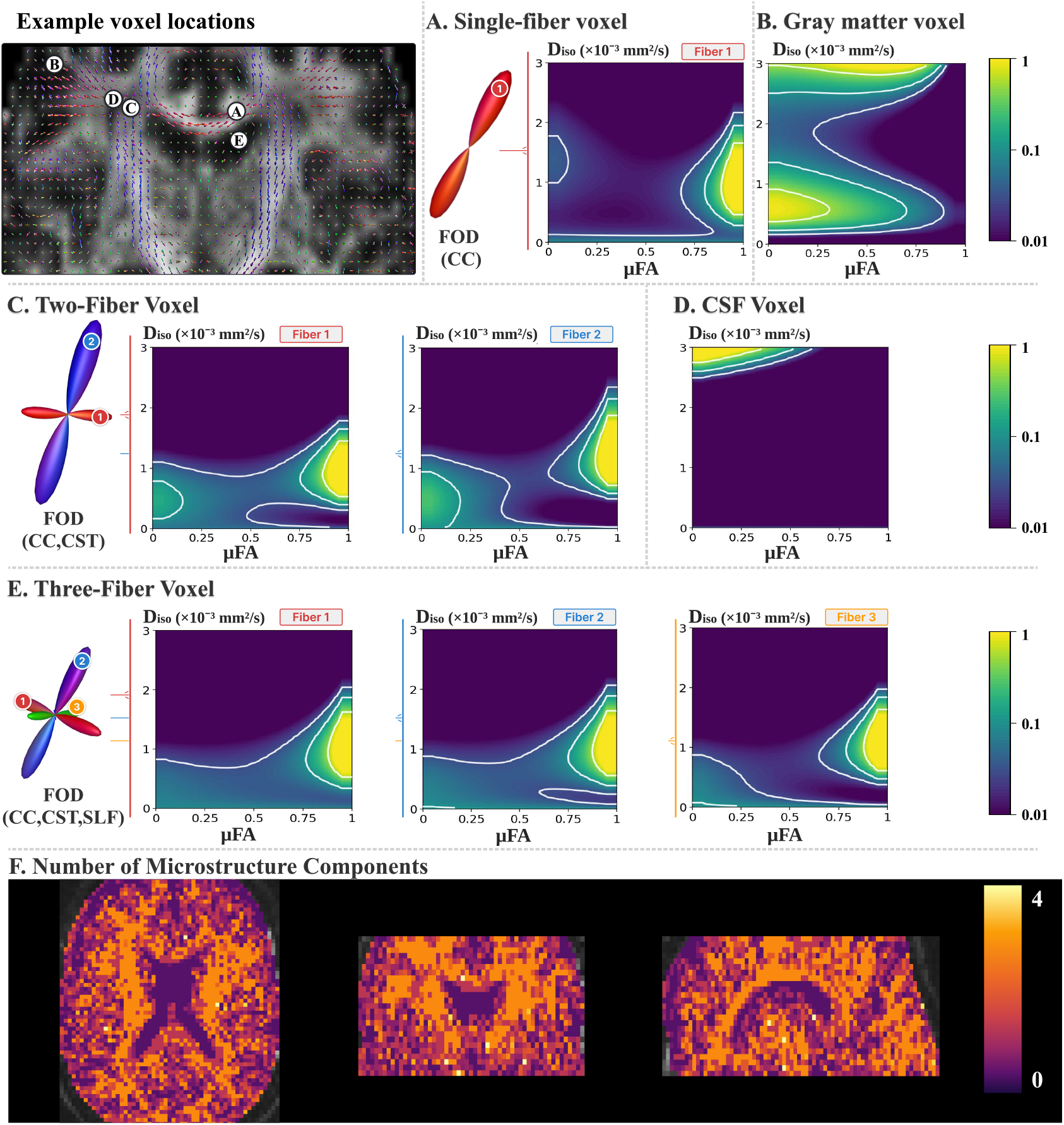
Fiber-specific microstructure by MaxEnt-DTD in representative in vivo voxels (B-tensor encoding). *Top left:* five voxels (A-E) marked on the FOD field (positions approximate). Each panel shows the FOD and the recovered distribution over *µ*FA and D_iso_; log-scaled density (a.u.) with white contours. For WM voxels (A, C, E) the per-fiber *P* (*µ*FA, D_iso_ | **u**) is shown (Fiber 1, red; 2, blue; 3, orange); for GM (B) and CSF (D) the voxel-level *P* (*µ*FA, D_iso_). (A) Single-fiber (CC); gray matter; (C) two-fiber (CC, CST); (D) C3S7F; (E) three-fiber (CC, CST, SLF). (F) Number of microstructure components per voxel, shown in representative axial, coronal, and sagittal slices.

Fig.11(B) shows the *µ*FA-D_iso_ distribution for a gray matter voxel, with two components corresponding to low and high D_iso_ values. The high-D_iso_ component, around 3 *µ*m^2^*/*ms, was similar to the component observed in the CSF voxel in Fig.11(D).

Fig.11(F) shows the average number of microstructure components, i.e., the peaks of the *µ*FA-D_iso_ distributions, in representative axial, coronal, and sagittal slices. Unlike the results in Fig.8, most WM voxels had a similar number of components, whereas the single-fiber voxels in Fig.8 had more components than other WM regions. GM voxels had 1 to 2 components and fewer components than WM voxels. The improved consistency in the number of components across WM regions demonstrates the benefit of B-tensor encoding for improving microstructure estimation. We note that the histogram as in Fig. 8(F) is not available since no anatomical image is provided in this dataset to compute the tissue labelmap.

#### 4.3.3. Rotation-invariant microstructure measures

Fig. 12 shows the estimated rotation-invarant metrics for the B-tensor encoded dataset. The *µ*FA and MK_A_ maps showed the most differences compared to results in Fig. 9. Specifically, the *µ*FA measures showed broader ranges values with more significant constrast between WM and GM. The MaxEnt-DTD also showed lower values in MK measures and with more relialble MK_A_ measures for all three methods. But the MD-dMRI showed slower values for the three MK measures which are consistent to the results in Fig. 6. The sgnificant difference between the Fig. 12 and Fig. 9 highlight the importance of using B-tensor encoding to improve microstructure estimation.

**Figure 12:**
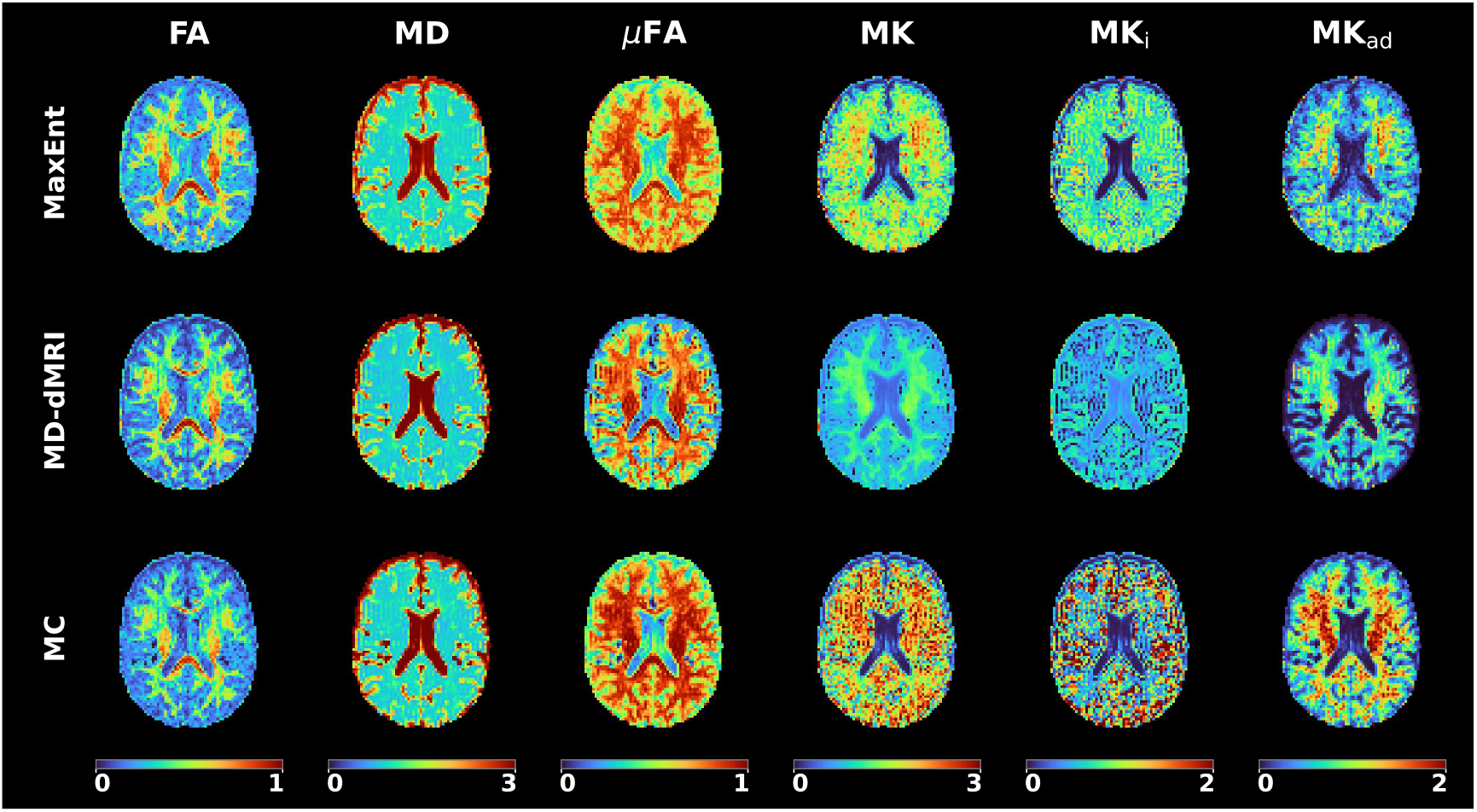
Rotation-invariant microstructure measures (B-tensor encoding). Rows: MaxEnt-DTD (MaxEnt), MD-dMRI, MC. Columns: fractional anisotropy (FA), mean diffusivity (MD, ×10^−3^ mm^2^*/*s), microscopic FA (*µ*FA), mean kurtosis (MK), and its isotropic and anisotropic parts (MK_i_, MK_ad_).

## 5. Discussion and Conclusion

This work introduces the MaxEnt-DTD framework for estimating diffusion tensor distributions (DTDs). The proposed method formulates DTD inversion as a convex optimization problem whose dimensionality is equal to the number of measurements, making the inversion tractable and scalable for whole-brain analysis. MaxEnt-DTD jointly characterizes fiber orientation distributions, orientation-dependent microstructure through conditional *µ*FA −D_iso_ distributions, and rotationally invariant metrics derived from the estimated cumulants. These complementary measures are obtained within a unified framework, which is not possible with conventional constrained spherical deconvolution (CSD) or parametric multi-compartment models that estimate fiber orientation and microstructure separately.

The performance of MaxEnt-DTD was evaluated using synthetic and *in vivo* dMRI data acquired with standard linear tensor encoding (LTE) and advanced B-tensor encoding protocols. In synthetic data, MaxEnt-DTD resolved fiber crossings with inter-fiber angles as small as 30^◦^, providing approximately 10^◦^ higher angular resolution than CSD-based methods. Conventional CSD methods assume that the diffusion signal can be modeled as a spherical convolution between a fixed fiber response function and an underlying FOD (Tournier et al., 2007; Jeurissen et al., 2014). This fixed response function constrains all fiber populations within a voxel to share identical microstructural properties and does not provide a probabilistic description of microstructural variability within or across fiber populations. Although the Monte Carlo approach achieved a high percentage of correct peak detections, it exhibited substantially larger angular errors, likely reflecting its greater sensitivity to measurement noise. The improved angular resolution of MaxEnt-DTD may be attributed to its independence from a fixed fiber response function which and allows for fiber-specific microstructure. Its improved reliability compared to the MC method arise from the explicit modeling of measurement noise, the convex optimization formulation that guarantees a globally optimal solution, and the low-dimensional representation that substantially reduces the parameter space.

The HCP protocol showed superior angular resolution compared with the B-tensor protocol, which may be related to the larger number of LTE measurements and higher b-values used in the HCP protocol.

The experimental results also showed that MaxEnt-DTD provided accurate microstructure estimation, including fiber-specific *µ*FA−D_iso_ distributions and rotationally invariant metrics. In synthetic data, MaxEnt-DTD achieved lower Wasserstein distances to the ground truth than the Monte Carlo method, especially for complex multi-fiber structures. The rotation-invariant measures showed that MaxEnt-DTD provided accurate and reliable estimation of cumulant-based metrics and more accurate MK estimation than the standard MD-dMRI method. Similar observations were reported by (Reymbaut et al., 2020), who demonstrated the MC inversion method provided more accurate microstructure measures than the cumulant-based MD-dMRI approach. Our results showed that MaxEnt-DTD produced more reliable estimates than the MC method in cases with relatively low SNR, especially for the *µ*FA and MK measures. Moreover, the comparison between the HCP and B-tensor encoding results highlights the importance of advanced B-tensor encoding for improving estimation accuracy, especially for the *µ*FA and MK_A_ measures.

The *in vivo* results demonstrate the feasibility of MaxEnt-DTD for whole-brain analysis. The method resolved crossing fiber bundles using both HCP and B-tensor protocols and enabled characterization of fiber-specific microstructural properties. In most white-matter regions, the estimated distributions exhibited one to two dominant modes, whereas gray-matter regions generally showed fewer distinct compartments. Crossing-fiber regions often exhibited fewer modes in the fiber-specific conditional *µ*FA−D_iso_ distributions, which may be related to the dependence of estimation accuracy on the number of fiber bundles, as shown in the synthetic data. The B-tensor encoding *in vivo* data also showed improved contrast and reliability in the *µ*FA and MK_A_ measures, consistent with observations from the synthetic data.

Finally, we note several limitations of the MaxEnt-DTD method. The proposed adaptive sampling approach depends on the accuracy of FOD estimation in the first step and high-resolution sampling in the second step. This makes the approach potentially sensitive to estimation errors in the first step. High-resolution sampling also limits computational efficiency. More efficient sampling methods in high-dimensional spaces, such as those in (Robert and Casella, 2004; Richard and Zhang, 2007; Bugallo et al., 2017), may further improve computational efficiency and help scale the approach to even higher-dimensional spaces.

In summary, the results demonstrate that MaxEnt-DTD provides a reliable framework for estimating diffusion tensor distributions. The proposed GPU-based implementation makes whole-brain analysis computationally feasible while providing a unified framework for simultaneous estimation of fiber orientation and microstructure. Both synthetic and *in vivo* experiments demonstrate robustness to measurement noise and improved accuracy relative to existing methods.

## References

Alexander, D.C., Dyrby, T.B., Nilsson, M., Zhang, H., 2019. Imaging brain microstructure with diffusion mri: practicality and applications. NMR in Biomedicine 32, e3841.

de Almeida Martins, J.P., Topgaard, D., 2016. Two-dimensional correlation of isotropic and directional diffusion using nmr. Physical review letters 116, 087601.

de Almeida Martins, J.P., Topgaard, D., 2018. Multidimensional correlation of nuclear relaxation rates and diffusion tensors for model-free investigations of heterogeneous anisotropic porous materials. Scientific Reports 8. doi:10.1038/s41598-018-19826-9.

Basser, P.J., Mattiello, J., LeBihan, D., 1994a. Estimation of the effective self-diffusion tensor from the NMR spin echo. Journal of Magnetic Resonance, Series B 103, 247–254. doi:10.1006/jmrb.1994.1037.

Basser, P.J., Mattiello, J., LeBihan, D., 1994b. Mr diffusion tensor spectroscopy and imaging. Biophysical journal 66, 259–267.

Basser, P.J., Pierpaoli, C., 1996. Microstructural and physiological features of tissues elucidated by quantitative-diffusion-tensor MRI. Journal of Magnetic Resonance, Series B 111, 209–219. doi:10.1006/jmrb.1996.0086.

Batou, A., Soize, C., 2013. Calculation of lagrange multipliers in the construction of maximum entropy distributions in high stochastic dimension. SIAM/ASA Journal on Uncertainty Quantification 1, 431–451.

Beaulieu, C., 2002. The basis of anisotropic water diffusion in the nervous system–a technical review. NMR in Biomedicine: An International Journal Devoted to the Development and Application of Magnetic Resonance In Vivo 15, 435–455.

Bugallo, M.F., Elvira, V., Martino, L., Luengo, D., Míguez, J., Djurić, P.M., 2017. Adaptive importance sampling: The past, the present, and the future. IEEE Signal Processing Magazine 34, 60–79. doi:10.1109/MSP.2017.2694771.

Callaghan, P.T., 2011. Translational Dynamics and Magnetic Resonance: Principles of Pulsed Gradient Spin Echo NMR. Oxford University Press. doi:10.1093/acprof:oso/9780199556984.001.0001.

Chouzenoux, E., Moussaoui, S., Idier, J., Mariette, F., 2010. Efficient maximum entropy reconstruction of nuclear magnetic resonance t1-t2 spectra. IEEE Transactions on Signal Processing 58, 6040–6051.

Christiaens, D., Veraart, J., Cordero-Grande, L., Price, A.N., Hutter, J., Hajnal, J.V., Tournier, J.D., 2020. On the need for bundle-specific microstructure kernels in diffusion mri. NeuroImage 208, 116460.

Coelho, S., Chen, J., Szczepankiewicz, F., Fieremans, E., Novikov, D.S., 2026. Geometry of the cumulant series in diffusion mri. Nature Communications 17, 4220.

Consagra, W., Ning, L., Rathi, Y., 2025. A deep learning approach to multi-fiber parameter estimation and uncertainty quantification in diffusion mri. Medical Image Analysis 102, 103537.

De Santis, S., Drakesmith, M., Bells, S., Assaf, Y., Jones, D.K., 2014. Why diffusion tensor mri does well only some of the time: variance and covariance of white matter tissue microstructure attributes in the living human brain. Neuroimage 89, 35–44.

Descoteaux, M., Deriche, R., Knosche, T.R., Anwander, A., 2008. Deterministic and probabilistic tractography based on complex fibre orientation distributions. IEEE transactions on medical imaging 28, 269–286.

Eriksson, S., Lasič, S., Nilsson, M., Westin, C.F., Topgaard, D., 2015. Nmr diffusion-encoding with axial symmetry and variable anisotropy: Distinguishing between prolate and oblate microscopic diffusion tensors with unknown orientation distribution. The Journal of chemical physics 142.

Essayed, W.I., Zhang, F., Unadkat, P., Cosgrove, G.R., Golby, A.J., O’Donnell, L.J., 2017. White matter tractography for neurosurgical planning: a topography-based review of the current state of the art. NeuroImage: Clinical 15, 659–672. doi:10.1016/j.nicl.2017.06.011.

Fischl, B., 2012. Freesurfer. NeuroImage 62, 774–781. doi:10.1016/j.neuroimage.2012.01.021.

Gorski, K.M., Hivon, E., Banday, A.J., Wandelt, B.D., Hansen, F.K., Reinecke, M., Bartelmann, M., 2005. Healpix: A framework for high-resolution discretization and fast analysis of data distributed on the sphere. The Astrophysical Journal 622, 759–771.

Gudbjartsson, H., Patz, S., 1995. The Rician distribution of noisy MRI data. Mag- netic resonance in medicine 34, 910–914.

Heaton, N.J., 2005. Multi-measurement nmr analysis based on maximum entropy. US Patent 6,960,913.

Jaynes, E.T., 1957. Information theory and statistical mechanics. Physical review 106, 620.

Jaynes, E.T., 1982. On the rationale of maximum-entropy methods. Proceedings of the IEEE 70, 939–952.

Jelescu, I.O., Budde, M.D., 2017. Design and validation of diffusion mri models of white matter. Frontiers in physics 5, 61.

Jelescu, I.O., Veraart, J., Fieremans, E., Novikov, D.S., 2016. Degeneracy in model parameter estimation for multi-compartmental diffusion in neuronal tissue. NMR in Biomedicine 29, 33–47.

Jensen, J.H., Helpern, J.A., Ramani, A., Lu, H., Kaczynski, K., 2005. Diffusional kurtosis imaging: the quantification of non-gaussian water diffusion by means of magnetic resonance imaging. Magnetic Resonance in Medicine: An Official Journal of the International Society for Magnetic Resonance in Medicine 53, 1432–1440.

Jeurissen, B., Descoteaux, M., Mori, S., Leemans, A., 2019. Diffusion mri fiber tractography of the brain. NMR in Biomedicine 32, e3785.

Jeurissen, B., Szczepankiewicz, F., 2021. Multi-tissue spherical decon-volution of tensor-valued diffusion MRI. NeuroImage 245, 118717. doi:10.1016/j.neuroimage.2021.118717.

Jeurissen, B., Tournier, J.D., Dhollander, T., Connelly, A., Sijbers, J., 2014. Multi-tissue constrained spherical deconvolution for improved analysis of multi-shell diffusion mri data. NeuroImage 103, 411–426.

Jian, B., Vemuri, B.C., Özarslan, E., Carney, P.R., Mareci, T.H., 2007. A novel tensor distribution model for the diffusion-weighted mr signal. NeuroImage 37, 164–176.

Karan, P., Reymbaut, A., Gilbert, G., Descoteaux, M., 2022. Bridging the gap between constrained spherical deconvolution and diffusional variance decomposition via tensor-valued diffusion mri. Medical image analysis 79, 102476.

Lampinen, B., Szczepankiewicz, F., Mårtensson, J., van Westen, D., Sundgren, P.C., Nilsson, M., 2017. Neurite density imaging versus imaging of microscopic anisotropy in diffusion mri: a model comparison using spherical tensor encoding. Neuroimage 147, 517–531.

Lasič, S., Szczepankiewicz, F., Eriksson, S., Nilsson, M., Topgaard, D., 2014a. Microanisotropy imaging: quantification of microscopic diffusion anisotropy and orientational order parameter by diffusion mri with magic-angle spinning of the q-vector. Frontiers in Physics 2, 11.

Lasič, S., Szczepankiewicz, F., Eriksson, S., Nilsson, M., Topgaard, D., 2014b. Microanisotropy imaging: quantification of microscopic diffusion anisotropy and orientational order parameter by diffusion mri with magic-angle spinning of the q-vector. Frontiers in Physics 2, 11.

Lawrence, I., Lin, K., 1989. A concordance correlation coefficient to evaluate reproducibility. Biometrics , 255–268.

Le Bihan, D., Breton, E., Lallemand, D., Grenier, P., Cabanis, E., Laval-Jeantet, M., 1986. Mr imaging of intravoxel incoherent motions: application to diffusion and perfusion in neurologic disorders. Radiology 161, 401–407.

Magdoom, K.N., Pajevic, S., Dario, G., Basser, P.J., 2021. A new framework for MR diffusion tensor distribution. Scientific Reports 11, 2766. doi:10.1038/s41598-021-81264-x.

Mohammad-Djafari, A., 2015. Entropy, information theory, information geometry and bayesian inference in data, signal and image processing and inverse problems. Entropy 17, 3989–4027.

Nilsson, M., Szczepankiewicz, F., Lampinen, B., Ahlgren, A., et al., 2018. An open-source framework for analysis of multidimensional diffusion mri data implemented in matlab, in: Proceedings of the International Society for Magnetic Resonance in Medicine (ISMRM).

Ning, L., 2023a. Maximum-entropy estimation of joint relaxation-diffusion distribution using multi-te diffusion mri, in: International Conference on Medical Image Computing and Computer-Assisted Intervention, Springer. pp. 439–448.

Ning, L., 2023b. Maximum-entropy estimation of joint relaxation-diffusion distribution using multi-te diffusion mri, in: International Conference on Medical Image Computing and Computer-Assisted Intervention, Springer. pp. 439–448.

Ning, L., Szczepankiewicz, F., Nilsson, M., Rathi, Y., Westin, C.F., 2021a. Probing tissue microstructure by diffusion skewness tensor imaging. Scientific Reports 11, 135.

Ning, L., Szczepankiewicz, F., Nilsson, M., Rathi, Y., Westin, C.F., 2021b. Probing tissue microstructure by diffusion skewness tensor imaging. Scientific Reports 11, 135.

Ning, L., Westin, C.F., Rathi, Y., 2015. Estimating diffusion propagator and its moments using directional radial basis functions. IEEE transactions on medical imaging 34, 2058–2078.

Novikov, D.S., Fieremans, E., Jespersen, S.N., Kiselev, V.G., 2019. Quantifying brain microstructure with diffusion mri: Theory and parameter estimation. NMR in Biomedicine 32, e3998.

Novikov, D.S., Veraart, J., Jelescu, I.O., Fieremans, E., 2018. Rotationally-invariant mapping of scalar and orientational metrics of neuronal microstructure with diffusion mri. NeuroImage 174, 518–538.

Özarslan, E., Koay, C.G., Shepherd, T.M., Komlosh, M.E., İrfanoğlu, M.O., Pierpaoli, C., Basser, P.J., 2013. Mean apparent propagator (map) mri: a novel diffusion imaging method for mapping tissue microstructure. NeuroImage 78, 16–32.

Palombo, M., Ianus, A., Guerreri, M., Nunes, D., Alexander, D.C., Shemesh, N., Zhang, H., 2020. Sandi: A compartment-based model for non-invasive apparent soma and neurite imaging by diffusion mri. Neuroimage 215, 116835.

Peyré, G., Cuturi, M., et al., 2019. Computational optimal transport: With applications to data science. Foundations and Trends® in Machine Learning 11, 355–607.

Phillips, S.J., Anderson, R.P., Schapire, R.E., 2006. Maximum entropy modeling of species geographic distributions. Ecological Modelling 190, 231–259. doi:10.1016/j.ecolmodel.2005.03.026.

Raffelt, D.A., Tournier, J.D., Smith, R.E., Vaughan, D.N., Jackson, G., Ridgway, G.R., Connelly, A., 2017. Investigating white matter fibre density and morphology using fixel-based analysis. Neuroimage 144, 58–73.

Reymbaut, A., 2020. Diffusion anisotropy and tensor-valued encoding. New Developments in NMR 24, 68–102. doi:10.1039/9781788019910-00068.

Reymbaut, A., Caron, A.V., Gilbert, G., Szczepankiewicz, F., Nilsson, M., Warfield, S.K., Descoteaux, M., Scherrer, B., 2021a. Magic diamond: Multi-fascicle diffusion compartment imaging with tensor distribution modeling and tensor-valued diffusion encoding. Medical Image Analysis 70, 101988.

Reymbaut, A., Critchley, J., Durighel, G., Sprenger, T., Sughrue, M., Bryskhe, K., Topgaard, D., 2021b. Toward nonparametric diffusion-characterization of crossing fibers in the human brain. Magnetic Resonance in Medicine 85, 2815–2827.

Reymbaut, A., Mezzani, P., de Almeida Martins, J.P., Topgaard, D., 2020. Accuracy and precision of statistical descriptors obtained from multidimensional diffusion signal inversion algorithms. NMR in Biomedicine 33, e4267. doi:10.1002/nbm.4267.

Richard, J.F., Zhang, W., 2007. Efficient high-dimensional importance sampling. Journal of Econometrics 141, 1385–1411. doi:10.1016/j.jeconom.2007.02.001.

Robert, C.P., Casella, G., 2004. Monte Carlo Statistical Methods. Springer Texts in Statistics. 2nd ed., Springer-Verlag, New York.

Rubner, Y., Tomasi, C., Guibas, L.J., 2000. The earth mover’s distance as a metric for image retrieval. International Journal of Computer Vision 40, 99–121.

Shannon, C.E., 1948. A mathematical theory of communication. The Bell system technical journal 27, 379–423.

Sijbers, J., Den Dekker, A.J., Scheunders, P., Van Dyck, D., 1998. Maximum-likelihood estimation of Rician distribution parameters. IEEE Transactions on Medical Imaging 17, 357–361.

Skilling, J., Bryan, R., 1984. Maximum entropy image reconstruction: general algorithm. Monthly notices of the royal astronomical society 211, 111–124.

Sotiropoulos, S.N., Jbabdi, S., Xu, J., Andersson, J.L.R., Moeller, S., Auer-bach, E.J., Glasser, M.F., Hernandez, M., Sapiro, G., Jenkinson, M., Fein-berg, D.A., Yacoub, E., Lenglet, C., Van Essen, D.C., Ugurbil, K., Behrens, T.E.J., WU-Minn HCP Consortium, 2013. Advances in diffusion MRI acquisi-tion and processing in the Human Connectome Project. NeuroImage 80, 125–143. doi:10.1016/j.neuroimage.2013.05.057.

Sporns, O., Tononi, G., Kötter, R., 2005. The human connectome: a structural description of the human brain. PLoS computational biology 1, e42.

Szczepankiewicz, F., Hoge, S., Westin, C.F., 2019. Linear, planar and spherical tensor-valued diffusion MRI data by free waveform encoding in healthy brain, wa-ter, oil and liquid crystals. Data in Brief 25, 104208. doi:10.1016/j.dib.2019.104208.

Topgaard, D., 2017. Multidimensional diffusion MRI. Journal of Magnetic Resonance 275, 98–113. doi:10.1016/j.jmr.2017.02.014.

Topgaard, D., 2019. Diffusion tensor distribution imaging. NMR in Biomedicine 32, e4066.

Tournier, J.D., Calamante, F., Connelly, A., 2007. Robust determination of the fibre orientation distribution in diffusion mri: non-negativity constrained super-resolved spherical deconvolution. Neuroimage 35, 1459–1472.

Tuch, D.S., Reese, T.G., Wiegell, M.R., Makris, N., Belliveau, J.W., Wedeen, V.J., 2002. High angular resolution diffusion imaging reveals intravoxel white matter fiber heterogeneity. Magnetic Resonance in Medicine: An Official Journal of the International Society for Magnetic Resonance in Medicine 48, 577–582.

Van Essen, D.C., Smith, S.M., Barch, D.M., Behrens, T.E., Yacoub, E., Ugurbil, K., Consortium, W.M.H., et al., 2013. The wu-minn human connectome project: an overview. Neuroimage 80, 62–79.

Villani, C., 2009. Optimal Transport: Old and New. volume 338 of Grundlehren der mathematischen Wissenschaften. Springer, Berlin, Heidelberg. doi:10.1007/978-3-540-71050-9.

Wedeen, V.J., Wang, R.P., Schmahmann, J.D., Benner, T., Tseng, W.Y.I., Dai, G., Pandya, D.N., Hagmann, P., D’Arceuil, H., de Crespigny, A.J., 2008. Diffusion spectrum magnetic resonance imaging (dsi) tractography of crossing fibers. Neuroimage 41, 1267–1277.

Westin, C.F., Knutsson, H., Pasternak, O., Szczepankiewicz, F., Özarslan, E., van Westen, D., Mattisson, C., Bogren, M., O’donnell, L.J., Kubicki, M., et al., 2016. Q-space trajectory imaging for multidimensional diffusion mri of the human brain. Neuroimage 135, 345–362.

Zhang, H., Schneider, T., Wheeler-Kingshott, C.A., Alexander, D.C., 2012. NODDI: practical in vivo neurite orientation dispersion and density imaging of the human brain. NeuroImage 61, 1000–1016. doi:10.1016/j.neuroimage.2012.03.072.

